# The conserved serine transporter SdaC moonlights to enable self recognition

**DOI:** 10.1101/2021.02.01.428846

**Authors:** Achala Chittor, Karine A. Gibbs

**Author notes:** To whom correspondence should be addressed via.

## Abstract

Cells can use self recognition to achieve cooperative behaviors. Self-recognition genes principally evolve in tandem with partner self-recognition alleles. However, other constraints on protein evolution could exist. Here, we have identified an interaction outside of self-recognition loci that could constrain the sequence variation of a self-recognition protein. We show that during collective swarm expansion in *Proteus mirabilis*, self-recognition signaling co-opts SdaC, a serine transporter. Serine uptake is crucial for bacterial survival and colonization. Single-residue variants of SdaC reveal that self recognition requires an open conformation of the protein; serine transport is dispensable. A distant ortholog from *Escherichia coli* is sufficient for self recognition; however, a homologous serine transporter, YhaO, is not. Thus, SdaC couples self recognition and serine transport, likely through a shared molecular interface. Understanding molecular and ecological constraints on self-recognition proteins can provide insights into the evolution of self recognition and emergent collective behaviors.

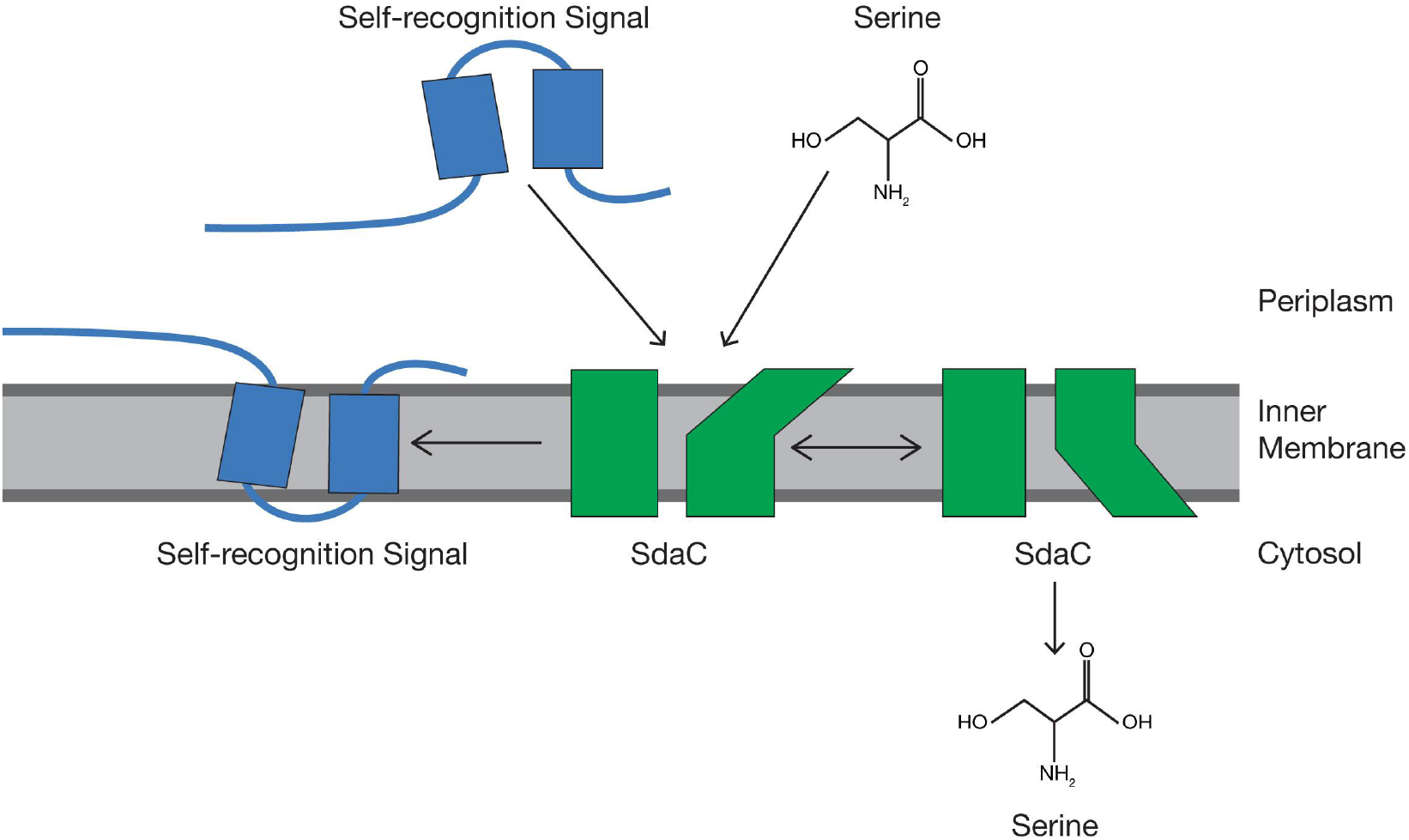

## Introduction

Self recognition regulates diverse processes in organisms across the tree of life. These vital roles include cell-cell communication, morphogenesis, and cooperation. For example, individual neurons express their identity through a unique set of clustered protocadherin proteins that interact between cells to inform neuronal self-avoidance and proper circuit formation (Kostadinov & Sanes, 2015; Lefebvre et al., 2012; Molumby et al., 2016). Kin cells of filamentous fungi can fuse to share resources in a syncytial lifestyle (Fischer & Glass, 2019; Gonçalves et al., 2020). And social microbes can identify, and coordinate with, kin during group migrations and fruiting body formation (Asfahl & Schuster, 2017; Gibbs et al., 2008; Gruenheit et al., 2017; Hirose et al., 2017; Pathak et al., 2013; Wenren et al., 2013). Self-recognition proteins often contain regions—stretches of amino acids—that vary between lineages to barcode a range of identities. The molecular mechanisms that regulate variation in self-recognition proteins are not fully elucidated.

From studies on eukaryotes, we know that protein-protein interactions affect how self-recognition proteins change over time. For clustered protocadherins, residues co-evolve across the homophilic interaction interface (Nicoludis et al., 2015, 2016, 2019). Allelic diversity of self-recognition genes results from balancing selection (Gruenheit et al., 2017; Noonan et al., 2003; Wu, 2005; Zhao et al., 2015). Still, no comprehensive model unifies genetics and molecular mechanisms. A similar gap between gene evolution and protein biochemistry exists for microbial self recognition. One can directly address this schism by studying the broader interaction networks that shape self-recognition evolution, both in individual microbial cells and populations.

Many microbes use self recognition to engage in collective behaviors selectively with kin. Microbial populations can build large, structured biofilms or migrate collectively across surfaces (Strassmann et al., 2011; West et al., 2007). For example, *P. mirabilis* cells engage in a collective migration known as swarming, allowing populations to cover surfaces efficiently. Individual cells communicate using self-recognition proteins to discern clonal siblings from others. Siblings gain preferred access to collective motility; non-self cells enter a transient altered state, resulting in exclusion from the swarming population (Cardarelli et al., 2015; Saak & Gibbs, 2016; Tipping & Gibbs, 2019). Unlike many cooperative self-recognition proteins found on the cell’s surface (Hirose et al., 2017; Pathak et al., 2013)*, P. mirabilis* cells inject the protein signal IdsD into adjacent neighbors. Mechanisms for localization and downstream signaling in the recipient cell are unknown.

For delivery into the recipient cell, IdsD may interact with a receptor protein residing in the inner membrane. Models propose that IdsD localizes to the inner membrane of recipient *P. mirabilis* cells (Cardarelli et al., 2015; Zepeda-Rivera et al., 2018), but there is less clarity about the delivery mechanism. A comparable model is the contact-dependent inhibition toxin, CdiA. In *Escherichia coli*, CdiA can promote collective behavior through cell-cell adhesion. Transferred from one cell into its neighbor, CdiA requires outer and inner membrane receptors for delivery to a recipient cell’s cytoplasm (Willett et al., 2015). Uptake depends on the interaction between CdiA’s modular translocation domain and a species-specific host receptor (Ruhe et al., 2013, 2017). None of IdsD’s species-specific binding partners—the chaperone IdsC in the donor cell and its self-recognition partner IdsE in the recipient cell (Cardarelli et al., 2015; Zepeda-Rivera et al., 2018)—could reasonably act as a receptor for delivery to the inner membrane. Although receptors on the inner or outer membrane have remained elusive, identifying these proteins is necessary to describe self-recognition pathways and elucidate further constraints on IdsD’s evolution.

With this in mind, consider bacteriophage-host interactions as a framework for studying the molecular evolution of self-recognition proteins. Many phage receptors are nutrient transporters on the outer and inner membranes. Phages exploit these host transporters, leading to rapid coevolution of competing proteins—an “arms race,” as reported for outer membrane proteins (de Jonge et al., 2019; Hampton et al., 2020). While mutations in the receptor can disrupt interactions and prevent killing, modifying these transporters can also carry fitness costs due to the importance of nutrient uptake (Mangalea & Duerkop, 2020; Meaden et al., 2015). Receptors for self-recognition proteins, particularly nutrient transporters, could likewise carry both costs and benefits. Discerning these interactions could reveal molecular constraints on the evolution of self-recognition systems.

Ids-mediated self recognition in *P. mirabilis* provides an opportunity to examine this concept. Cells can bypass recognition-triggered limitations by removing the self-recognition genes or the cell-to-cell transport system (Saak & Gibbs, 2016; Zepeda-Rivera et al., 2018). Here we show that another mechanism is the disruption of the serine transporter SdaC. Self-recognition signaling specifically uses SdaC during collective motility. By analyzing single-residue variants to alter protein state, we show self-recognition signaling requires an open conformation but not serine transport. Supporting the importance of a protein interface, we saw that an ortholog from *E. coli* is sufficient for self recognition. A related serine transporter in *P. mirabilis*, YhaO, is not. Therefore, SdaC moonlights to couple self recognition with serine transport, revealing a critical interaction and a potential regulator for self-recognition protein evolution.

## Results

Organisms can escape self-recognition control using several strategies. Spontaneous mutations that restore collective motility have removed self-recognition communication, either by disrupting self-recognition genes themselves or disrupting the necessary transport system (Saak & Gibbs, 2016). To identify additional mechanisms, we modified an unbiased assay for isolating self-recognition escape mutants emerging from an initial swarm colony [(Figure 1A), (Saak & Gibbs, 2016)]. We selected twelve novel independent mutant strains for further study. Whole-genome sequencing showed that these strains contained mutations in the gene *sdaC*; all, except two, were deleterious to a full-length protein (Figure 1B). We reasoned that *sdaC* disruption might constitute another mode of escape from recognition-based swarm exclusion.

**Figure 1:**
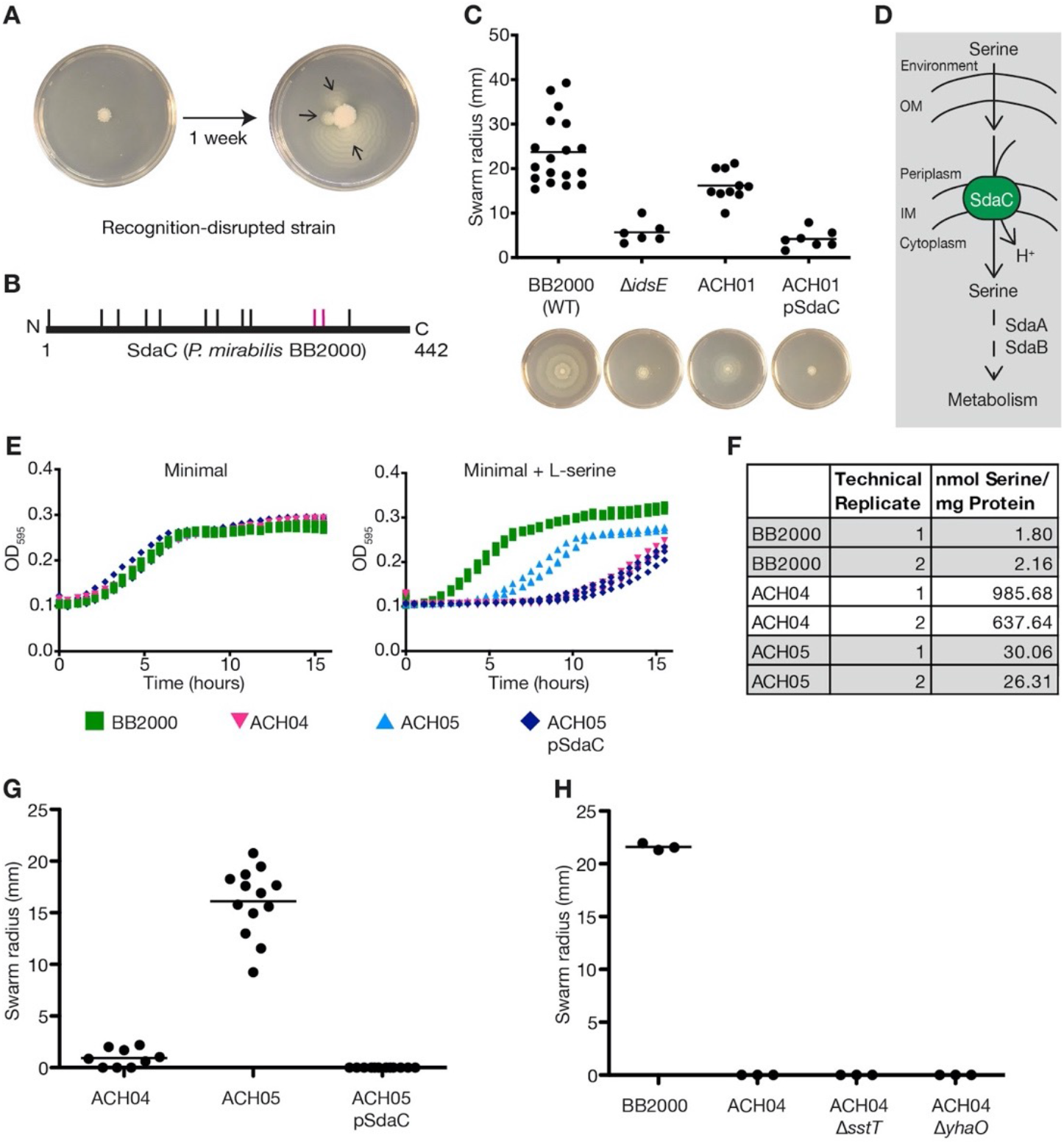
SdaC is required for Ids-mediated self recognition and is the dominant serine transporter during swarming. A) Strains containing *de novo* mutations emerge (marked with arrows) from a restricted swarm colony of strain BB2000 Δ*ids* pIdsBBΔ*idsE* (Saak & Gibbs, 2016). Images of swarm plate at the end of standard two day incubation (left) and after incubating an additional one week (right). B) Isolated mutations occur throughout SdaC protein sequence; relative location denoted by tick marks. Magenta lines mark missense mutations. Black lines denote nonsense mutations. See Table S1 for specific sites. C) Swarm radius measured from swarm assay of BB2000 empty vector, Δ*idsE* empty vector, ACH01 [BB2000 Δ(*idsE*, *sdaC*)] empty vector, and ACH01 pSdaC. Representative plate images shown below. D) Brief prediction of serine uptake and utilization pathway in *P. mirabilis*, which was constructed based on the homologies of SdaA, SdaB, and SdaC to *E. coli* proteins. E) Growth curve of BB2000 empty vector, ACH04 [BB2000 Δ(*sdaA*, *sdaB*)] empty vector, ACH05 [BB2000 Δ(*sdaA*, *sdaB*-*sdaC*)] empty vector, and ACH05 pSdaC in minimal medium (left) and minimal medium plus 10mM L-serine (right) for three biological replicates. F) Serine concentration calculated from LC-MS results of total swarmer cell lysates of BB2000, ACH04, and ACH05 and normalized to total protein content for two technical replicates. G) Swarm radius measured (from edge of inoculum) from swarm assay of ACH04 empty vector, ACH05 empty vector, and ACH05 pSdaC. CFUs per swarm colony shown in Figure S2E. Representative plate images in Figure S4. H) Swarm radius measured from swarm assay of strains BB2000, ACH04, ACH04 Δ*sstT*, ACH04 Δ*yhaO*.

Direct deletion and complementation of *sdaC* in a clean genetic background could confirm this proposed role. We generated a non-polar deletion of *sdaC* in the non-self Δ*idsE* background (Zepeda-Rivera et al., 2018), resulting in strain ACH01. In a colony of the Δ*idsE* strain, each cell perceives others as non-self, resulting in restricted collective swarm motility (Figure 1C). Strain ACH01 showed a relative three-fold increase in collective swarm motility. By contrast, plasmid-based expression of *sdaC* in ACH01 restricted motility back to the non-self strain’s level (Figure 1C). The rescue of collective motility in ACH01 is not due to a change in the self-recognition protein’s transport. The *sdaC* gene was dispensable for IdsD secretion as measured using a self-recognition assay based on boundary formation between colliding swarming populations (Figure S1). Thus, *sdaC* is necessary for self-recognition signal transduction in the receiving cell.

The SdaC protein is a predicted membrane-bound serine transporter but without confirmed function in *Proteus* spp. In *E. coli* and other enteric Gram-negative bacteria, SdaC is an integral inner membrane protein that brings serine into the cell coupled to a proton (Hama et al., 1988; Shao et al., 1994; Velayudhan et al., 2004). Serine deaminases (SdaA and SdaB) primarily metabolize imported serine to pyruvate (Shao & Newman, 1993; Su et al., 1989; Velayudhan et al., 2004)*. Proteus* spp. contain genes with sequence similarity to these serine import and utilization genes (Figure 1D). We confirmed membrane localization of *P. mirabilis* SdaC by expressing a mCherry fusion from its native promoter (Figure S3). As with *E. coli* (Baba et al., 2006), deletion of *sdaC* does not produce any significant growth or motility defects in *P. mirabilis* (Figure S2A-B). Serine uptake and utilization in *P. mirabilis* via SdaA, SdaB, and SdaC resemble *E. coli* (Figure 1D). Therefore, we can use what is known in *E. coli* to develop tools to investigate the function of SdaC further.

Serine transport should contribute to the internal serine pool and could shift growth dynamics. Deleting the *E. coli* serine deaminases increases internal serine concentration, causing physiological effects, including growth defects in minimal medium and cell wall instability (Hama et al., 1990, 1991; X. Zhang et al., 2010; X. Zhang & Newman, 2008). Informed by these results, we made a *P. mirabilis* strain with a serine-dependent reporter phenotype. Removing the comparable serine deaminases (the *sdaA* and *sdaB* genes) resulted in strain ACH04 (Figure S2C). Consistent with serine toxicity observed in *E. coli*, this strain had the expected delayed growth in a minimal medium containing exogenous serine but not in the absence of serine (Figure 1E). Swarm colony cells were also lysed and subjected to LC-MS analysis to measure serine concentrations. ACH04 cells contained about 400-fold higher intracellular serine than the wild-type parent (Figure 1F). They also displayed less swarm expansion than wildtype (Figure 1G). Based on subsequent experiments (Figure S2D-G) and earlier reports (Little et al., 2019; X. Zhang et al., 2010), serine-induced cell wall instability in strain ACH04 probably causes the reduced growth rate and swarm colony expansion in nutrient-rich conditions. Nonetheless, swarm expansion in the ACH04 strain background can serve as a readout for internal serine levels.

We reasoned that if SdaC were a serine transporter in *P. mirabilis*, then deletion of *sdaC* would alleviate defects in strain ACH04. ACH05, the engineered strain containing a *sdaC* deletion in the ACH04 background, showed a partial rescue of growth in a minimal medium with serine (Figure 1E). Internal serine concentrations decreased by ~ 30-fold compared to ACH04 (Figure 1F), while swarm expansion increased by roughly 8-fold (Figure 1G). Adding back SdaC to ACH05, through plasmid-based expression, reproduced the growth and colony expansion defects (Figure 1E, G). However, two other predicted serine transporters, *sstT* and *yhaO* (Connolly et al., 2016; Ogawa et al., 1998), could play a role. To examine any potential contributions to internal serine during swarming, we individually removed *sstT* and *yhaO* from the ACH04 background. Neither strain showed swarm colony expansion and instead looked like the ACH04 parental strain (Figure 1H, S5). Thus, SdaC is indeed a serine transporter in *P. mirabilis* and is dominant during collective motility.

Of the possible ways in which SdaC could function in self-recognition, two seemed most probable. SdaC serine transport could regulate downstream self-recognition signaling by modulating internal serine levels. Alternatively, a specific conformation of SdaC could be a required binding interface. Indeed, point mutations in SdaC already hinted at a molecular mechanism for its function in signal transduction. Two independent full-length SdaC disruptions emerged from the suppressor screen: G328V and G332R (Figure 1B). These residues sit in a predicted interface that stabilizes the open conformation in LeuT, a similar transport protein [Figure 2A, (Krishnamurthy & Gouaux, 2012)]. In combination with the assay toolkit, additional reduced-function point mutations could reveal SdaC’s role in self recognition.

**Figure 2:**
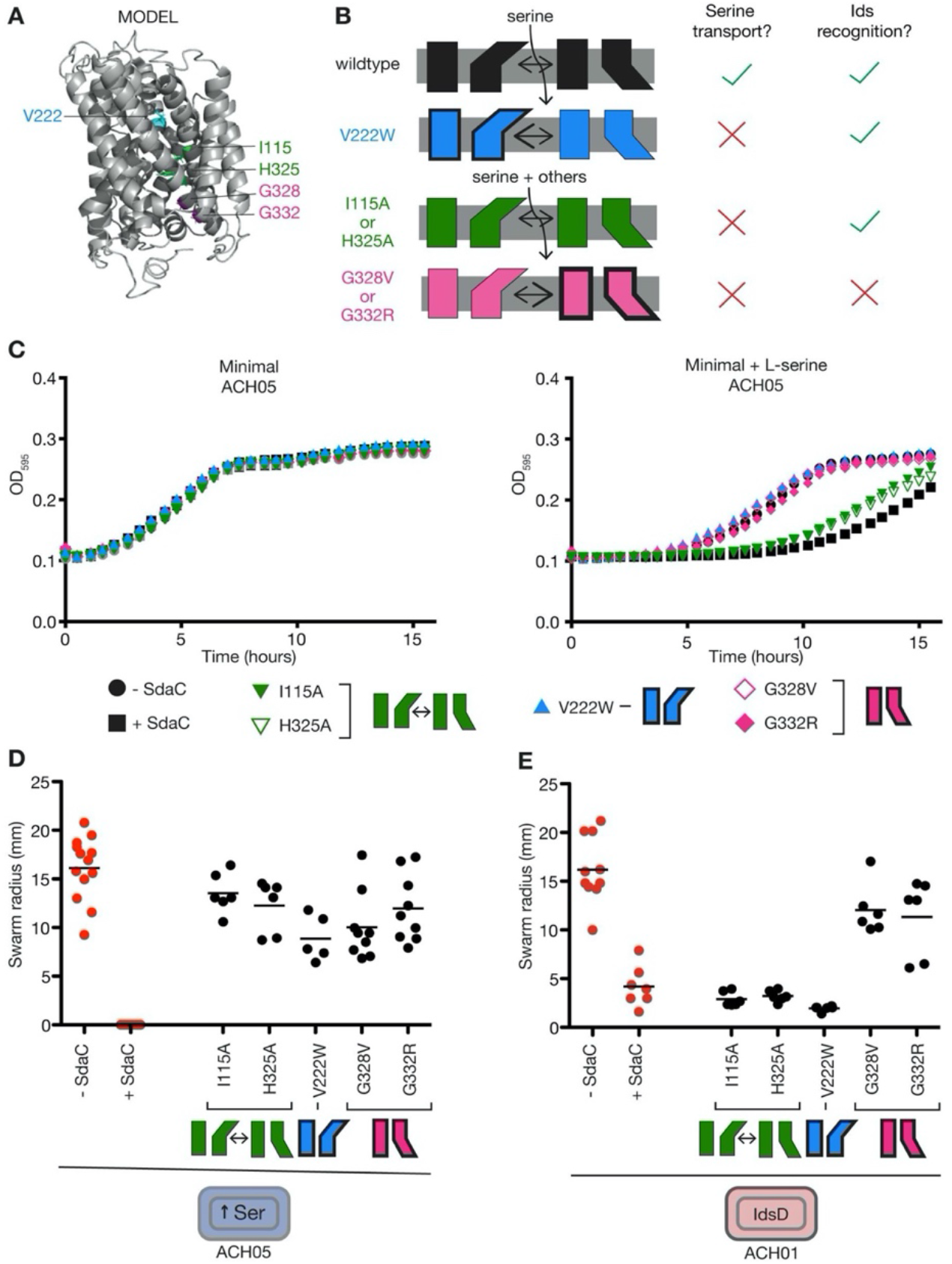
SdaC’s conformation determines serine transport and IdsD signaling. A) PyMol figures of a structural model for *P. mirabilis* SdaC from I-TASSER (C-score = −0.82, TM-score = 0.61±0.14, RMSD = 8.9±4.6 Å). Targeted residues are labeled. B) Predicted conformational bias and serine transport changes are shown as a protein cartoon for each mutation: V222W is open-biased, I115A and H325A are non-specific in substrate affinity, G328R and G332V are closed-biased. Summary of (B-E) results are shown on the right with a check (success) or cross (failure). C) Growth curves of each strain, expressed from the native promoter in the ACH05 [BB2000 Δ(*sdaA*, *sdaB-sdaC*)] strain background, in minimal medium (left) or minimal medium plus 10 mM L-serine (right) for three biological replicates. ACH05 empty vector and ACH05 pSdaC data were copied from Figure 1E and colored in red for comparison. D) Swarm radius measured from assay of same strains from (C). ACH05 empty vector and ACH05 pSdaC are copied from Figure 1G and colored in red for comparison. Blue cell icon indicates that intracellular serine levels are high in the ACH05 background when SdaC is active, leading to restricted swarm expansion. E) Swarm radius measured from swarm assay of each mutant expressed from the native promoter in the ACH01 [BB2000 Δ(*idsE*, *sdaC*)] strain background. ACH01 empty vector and ACH01 pSdaC are copied from Figure 1C and colored in red for comparison. Red cell icon indicates that IdsD and downstream recognition signaling are active in the ACH01 background when SdaC is active, leading to restricted swarm expansion.

A structural model for SdaC resembles LeuT-fold proteins, providing a template for generating variants with biased conformations. We used the structure prediction program I-TASSER (Roy et al., 2010; Yang et al., 2015; Y. Zhang, 2008) to make this model based on the primary amino acid sequence and solved structures of similar proteins (Figure 2A). LeuT-fold transporters sample at least three conformations: open (outward-facing), closed (inward-facing), and an intermediate state (Bozzi et al., 2019; Krishnamurthy & Gouaux, 2012). A V222W mutation would reasonably bias the SdaC protein to the open conformation (Figure 2A-B); an equivalent conversion in NRAMP favors the open conformation (Bozzi et al., 2019)]. The mutations in the suppressor strains, G328V and G332R, are predicted to bias SdaC to the closed conformation [Figure 2A-B, (Krishnamurthy & Gouaux, 2012)]. Altering either of two residues (I115A and H325A) in the substrate-binding site would allow non-specific transport of additional amino acids (Figure 2A-B) while continuing to sample both open and closed conformations. Each mutation was introduced independently into *sdaC* expressed from its native promoter on a plasmid. The engineered variants were visualized using an N-terminal mCherry fusion comparable to the wildtype and localized to the *P. mirabilis* cell envelope (Figure S3). The mutant strains provide molecular levers to distinguish between contributions to serine transport versus self recognition.

Bias toward an open or closed conformation should restrict serine transport compared to that of the wildtype and the mutant that transports non-specifically. Therefore, we tested transport function in the ACH05 background, which lacks the serine deaminases. All strains grew equivalently in minimal medium (Figure 2C). As shown earlier, ACH05 has lowered internal serine and grows like wild-type in a minimal medium with exogenous serine (Figure 1E). The addition of transgenic SdaC resulted in attenuated growth (Figure 1E). The open-biased (V222W) and closed-biased (G328V, G332R) variants grew like ACH05 in minimal medium plus excess serine (Figure 2C). However, the non-specific variants (I115A, H325A) showed attenuated growth in minimal medium plus excess serine, much like ACH05 pSdaC (Figure 2C). This reduced liquid growth did not translate to altered swarm expansion. All mutant strains expanded beyond a radius of 10 mm, similar to ACH05, instead of being restricted as observed in ACH05 pSdaC (Figure 2D). These results support that serine transport is not itself required for swarm colony expansion under these conditions.

Suppose self-recognition relies on SdaC-mediated serine transport. In that case, the conformations biased to open or closed should prevent Ids-mediated recognition signaling, allowing swarm colonies of BB2000 Δ*idsE* (all non-self cells) to expand. To test this hypothesis, we introduced each SdaC variant into the ACH01 background (Figure 1C). The closed-biased variants (G328V and G332R) showed increased swarm colony expansion (Figure 2E), consistent with these mutations emerging from the original suppressor screen (Figure 1A-B). By contrast, the open-biased (V222W) and non-specific variants (I115A and H325A) exhibited restricted swarm expansion comparable to that of ACH01 pSdaC, which is the wild-type protein (Figure 2E). Altogether, these results indicate that self-recognition signaling requires for SdaC to sample the open conformation, but not transport serine. These SdaC functions are distinct and overlapping.

The natural sequence variation among similar proteins provides an avenue for understanding what molecular aspects of SdaC might be critical for self recognition. SdaC from *E. coli* (SdaC-Ecol) is a diverged ortholog whose function is understood (Shao et al., 1994). As stated earlier, SdaC is in a broader conserved pathway with SdaA and SdaB, apparently shared between *E. coli* and *P. mirabilis.* On the other hand, YhaO, another predicted serine/H+ symporter in *P. mirabilis*, is a distant homolog. In *E. coli*, YhaO transports primarily D-serine, and to a lesser degree, L-serine (Connolly et al., 2016). YhaO (*P. mirabilis*) shares much less sequence identity with SdaC-Pmir than SdaC-Ecol; yet, conserved residues are visible throughout the protein, especially in the predicted transmembrane domains (Figure 3). I-TASSER-based predictions for the structures of SdaC-Ecol and YhaO also have a LeuT fold and share the same I-TASSER templates as SdaC-Pmir (Figure S7). SdaC-Pmir, SdaC-Ecol, and YhaO share sequence similarities and predicted tertiary structures.

**Figure 3:**
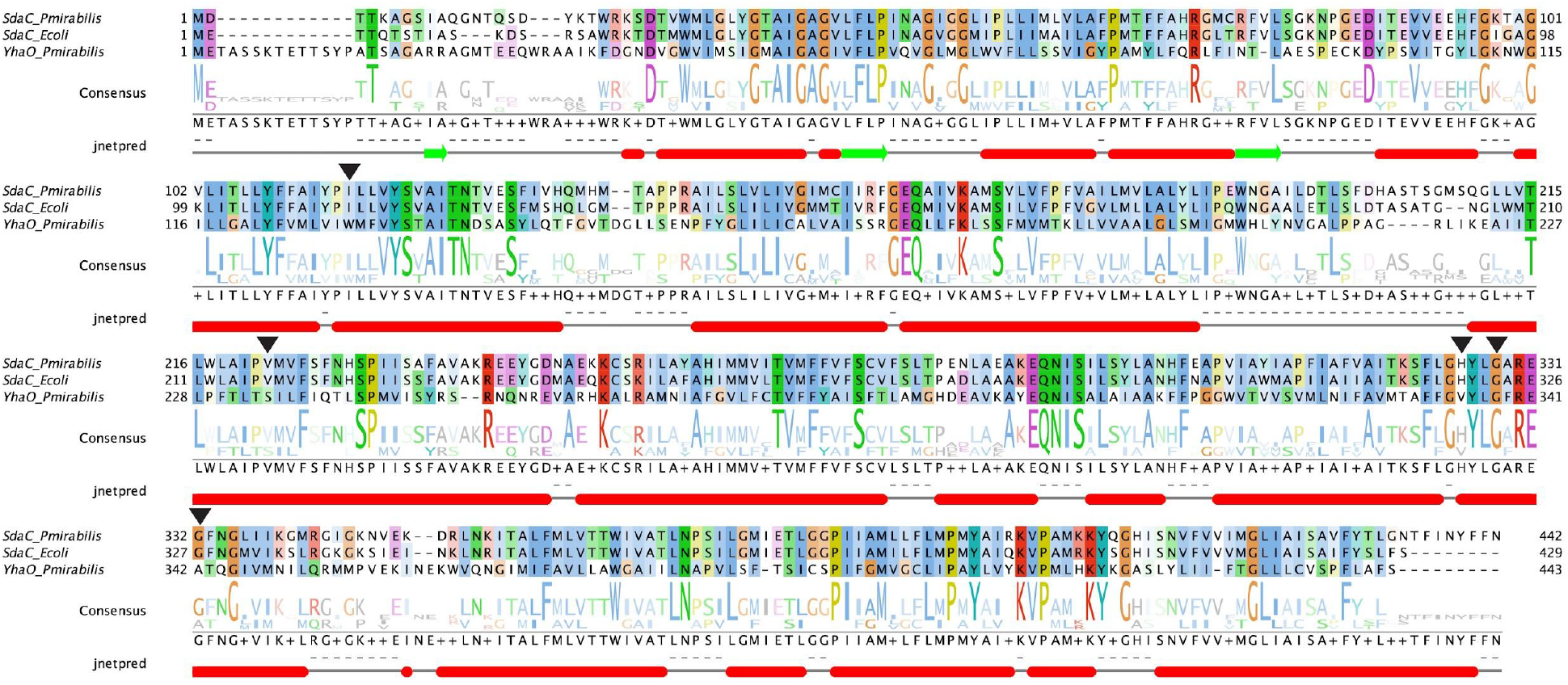
Sequence alignment of SdaC orthologs from *P. mirabilis* and *E. coli* and the homolog YhaO from *P. mirabilis*. Aligned to each other are SdaC from *P. mirabilis* BB2000 (BB2000_0742), SdaC from *E. coli* K-12 MG1655, and YhaO from BB2000 (BB2000_2747) using Clustal Omega (Madeira et al., 2019; Sievers et al., 2011). The alignment was modified in Jalview (Waterhouse et al., 2009) to show the consensus sequence and logo plot as well as secondary structure prediction using JPred (red ovals for alpha-helices and green arrows for beta-sheets). Targeted residues are labeled with black arrowheads.

Suppose sampling an open conformation is the critical molecular mechanism for SdaC’s function in self recognition. In that case, SdaC-Ecol and YhaO should be able to replace SdaC in *P. mirabilis*. To interrogate this hypothesis, we expressed SdaC-Ecol and YhaO from an inducible promoter in the ACH05 background. The resultant strains showed growth defects in minimal medium with added serine but not in minimal medium (Figure 4A). Therefore, both SdaC-Ecol and YhaO are sufficient to substitute for SdaC’s serine transport function in *P. mirabilis*. Next, SdaC-Ecol and YhaO were expressed from an inducible promoter in the non-self ACH01 background and subjected to the swarm expansion assay. Strains producing SdaC-Pmir or SdaC-Ecol showed similarly restricted swarm expansion (Figure 4B). The strain producing YhaO expanded more fully (Figure 4B). Despite the sequence divergence, only SdaC-Ecol could substitute for SdaC and reproduce self recognition.

**Figure 4:**
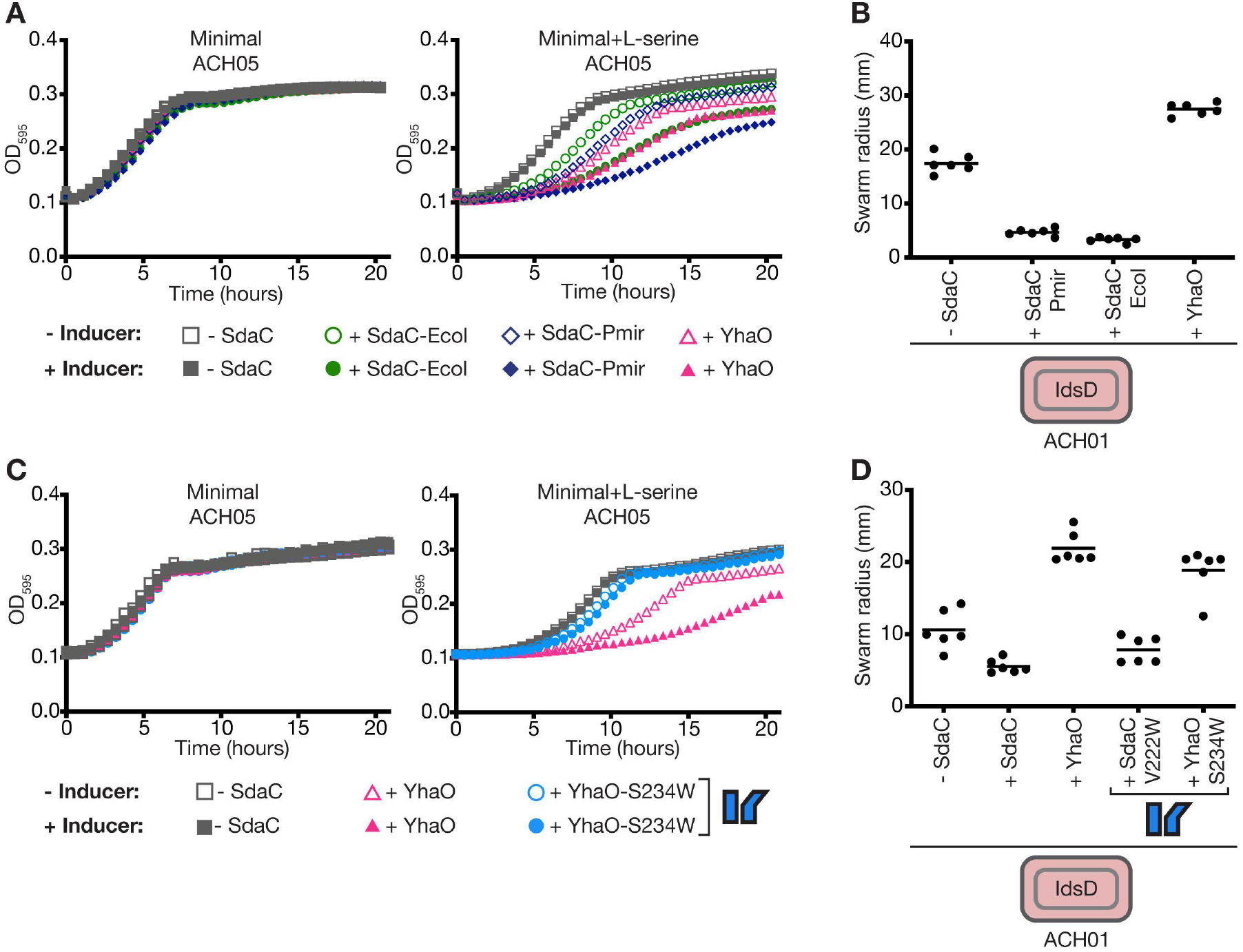
SdaC homologs share a similar predicted structure and function but only the SdaC ortholog from *E. coli* can complement in self recognition. A) Growth curve of ACH05 [BB2000 Δ(*sdaA, sdaB-sdaC*)] expressing an empty vector, pTet-SdaC-Pmir, pTet-SdaC-Ecol, or pTet-YhaO without inducer (empty icons) or with inducer (filled icons) in minimal medium (left) and minimal medium plus 10mM L-serine (right). Mean of six biological replicates shown. B) Swarm radius measured from swarm assay of ACH01 [BB2000 Δ(*idsE*, *sdaC)*] carrying empty vector, pTet-SdaC-Pmir, pTet-SdaC-Ecol, or pTet-YhaO. Red cell icon indicates that IdsD and downstream recognition signaling are active in the ACH01 strain background when SdaC is active, leading to restricted swarm expansion. C) Growth curve of ACH05 expressing an empty vector, pTet-YhaO, or pTet-YhaO-S234W (open-biased) with inducer (filled icons) or without inducer (empty icons) in minimal medium (left) and minimal medium plus 10mM L-serine (right). Mean of three biological replicates shown. D) Swarm radius measured from swarm assay of ACH01 carrying empty vector, pTet-SdaC-Pmir, pTet-SdaC-Pmir-V222W (open-biased), pTet-YhaO, or pTet-YhaO-S234W (open-biased). Red cell icon indicates that IdsD and downstream recognition signaling is active in the ACH01 strain background when SdaC is active, leading to restricted swarm expansion.

Theoretically, YhaO might not sufficiently sample an open conformation, so we introduced a point mutation to bias the protein. We constructed an open-biased S234W variant of YhaO analogous to the V222W variant of SdaC. Unlike wild-type YhaO, there was no growth defect for the open-biased variant expressed in ACH05 when grown in a minimal medium with serine (Figure 4C). This disrupted serine transport is consistent with the results of the open-biased SdaC variant (Figure 2C). We then moved the open-biased variant into the ACH01 background. The resultant strain showed swarm expansion similar to wild-type YhaO (Figure 4D). YhaO in the open conformation is sufficiently different from SdaC that it does not work in self recognition (Figure 4D). While SdaC’s transport function is conserved among similar proteins, there is specificity to its conformation and sequence needed for self-recognition signal transduction.

## Discussion

SdaC, a serine transporter, moonlights in the self-recognition signaling pathway. The transduced self-identity signal, IdsD, is a type VI secretion substrate predicted to localize to a recipient cell’s inner membrane (Cardarelli et al., 2015; Zepeda-Rivera et al., 2018). Though inner membrane transporters are not known as receptors for type VI secretion substrates, we propose that SdaC functions as a receptor to promote IdsD insertion. Consistent with this hypothesis, disrupting SdaC also provides resistance to microcin V and phage C1 (Gérard et al., 2005; Likhacheva et al., 1996). Multiple inner membrane proteins are nutrient transporters and receptors for incoming proteins. For example, the contact-dependent inhibition protein CdiA requires specific inner membrane transporters for proposed insertion (Ruhe et al., 2017; Willett et al., 2015). Based on work in CdiA, a predominant hypothesis is that the membrane protein’s functions work independently: nutrient transport versus translocation of protein. Our data expands this prior model. For SdaC, self recognition and nutrient transport are interdependent, likely affecting the protein’s evolutionary trajectory.

Cells need SdaC to sample an open conformation for either self recognition or serine transport to occur. Like CdiA receptors (Willett et al., 2015), self-recognition signaling and collective motility do not specifically require serine transport (Figure 2D-E). Removing SdaC allows cells to bypass Ids-mediated self recognition (Figure 1C). Non-self populations regained collective motility when SdaC was not functional (Figure 1A-C, Figure 2E). Mutations that biased SdaC to an open conformation were sufficient to permit Ids-mediated self recognition regardless of serine transport (Figure 2E). However, the open conformation is only accessible when SdaC undergoes the conformational dynamics needed for active transport. These two functions of SdaC are distinct but not independent.

High internal serine concentrations are toxic, inducing cell envelope stress and fitness defects of slower growth and no swarming (Figures 1E-G, 2C-E, S2D-G). Stopping serine transport by deleting SdaC rescues serine toxicity in mutant strains lacking serine deaminases (Figure 1E-G). However, deleting the two other serine transporters, SstT and YhaO, does not relieve serine toxicity during swarming (Figure 1H). Therefore, internal serine levels, partially controlled by SdaC activity, are important during collective motility. And it is during this collective motility that self recognition occurs (Tipping & Gibbs, 2019). Expanding upon proposals for phage receptors, the coupling of SdaC functions may limit the emergence of mutations in both pathways.

Ecological context could constrain SdaC evolution. In bacteriocin and phage receptors, the local ecology and associated fitness trade-offs impact the emergence of intersectional mutations (Feldgarden & Riley, 1999; Inglis et al., 2016; Mangalea & Duerkop, 2020; Meaden et al., 2015). Serine is a crucial metabolite for urinary tract and gut pathogens (Barroso-Batista et al., 2020; Brauer et al., 2019; Connolly et al., 2016; Kitamoto et al., 2019; Velayudhan et al., 2004). Moreover, serine homeostasis is vital. Elevated internal serine poisons cells, leading to growth defects and susceptibility to cell envelope stress [Figure S2D-G, (X. Zhang et al., 2010; X. Zhang & Newman, 2008)]. Too little serine starves cells of a significant amino acid. SdaC is the critical serine transporter during swarming (Figure 1E-H), a behavior correlated with disease (Armbruster et al., 2018; Kearns, 2010; Schaffer & Pearson, 2015). Further, the SdaA, SdaB, and SdaC serine uptake pathway holds for distantly related species, suggesting evolutionary conservation and importance (Figure 3, 4A). Our results support that SdaC is a bifunctional conserved molecular interface, constrained by *Proteus*’ ecology.

SdaC structure may limit plasticity and exploration of sequence space in buried regions. Often, exposed loops of receptor proteins are the main interaction interface based on outer membrane proteins co-opted by phages and toxins (Chatterjee & Rothenberg, 2012; Kleanthous, 2010; Ruhe et al., 2013). However, this model does not appear to hold for SdaC from *E. coli* and *P. mirabilis*. Sequences for the exposed periplasmic loops are ~ 65% identical (Figure 3). At the same time, the transmembrane regions are ~ 86% identical (Figure 3). Yet, *E. coli* SdaC can function in self recognition (Figure 4B) and transport (Figure 4A). By contrast, the *P. mirabilis* YhaO protein also aligns with SdaC (Figure 3) and transports serine (Figure 4A). However, YhaO cannot substitute for self recognition, even when biased into an open conformation (Figure 4B-D). Our data suggest that the membrane-localized pocket, predicted to be exposed conditionally in the open conformation, is the interaction interface (Figure 2). SdaC may follow conventional evolution ideas for integral membrane helical proteins, which are predicted to evolve slowly in buried regions due to molecular constraints (Oberai et al., 2009). Multi-conformation proteins are further constrained (Sharir-Ivry & Xia, 2017). The synergy between ecological fitness and structural constraints could slow the rate of SdaC sequence changes.

SdaC conservation can potentially regulate IdsD sequence drift to preserve signal fidelity. We have discussed the constraints on SdaC, but what constraints exist for the identity signal, IdsD? Self-recognition genes contain polymorphic regions (self-identity barcodes) that can serve as a proxy for relatedness (de Oliveira et al., 2019). A dominant model is that evolution of self-recognition proteins is constrained by interactions with themselves or other self-recognition proteins, driving variation in their interaction interface (Cardarelli et al., 2015; Hirose et al., 2017; Pathak et al., 2013). IdsD does bind other self-recognition proteins such as its chaperone IdsC (Zepeda-Rivera et al., 2018) and its partner recognition protein IdsE (Cardarelli et al., 2015). Extending this model, we hypothesize that SdaC, which is not a recognition protein, acts as an additional source of selective pressure. IdsD requires SdaC in a sequence-specific manner. Although predicted to have a similar structure and function, YhaO cannot replace SdaC for recognition (Figure 4). For uptake, IdsD must retain compatibility with SdaC. Therefore, SdaC potentially acts as a bottleneck for signal transduction in the receiving cell.

Molecular crosstalk mirrors observed interactions between nutrient availability, collective behavior, and self recognition in many organisms. Collective behaviors are associated with nutrient limitation in other microbes (Kundert & Shaulsky, 2019; Wall, 2014), fungi (Gonçalves et al., 2020), and plants (Palmer et al., 2016). Collective behaviors can allow for sharing of nutrients and promote developmental processes such as fruiting body formation. Self-recognition signaling allows preferential collective action with kin, an advantage during nutrient limitation. Functional coupling between self recognition and organism-relevant pathways such as nutrient transport, as shown for SdaC, could constrain identity signal evolution. For collective behaviors, nutrient uptake is a crucial regulator. Still, for other contexts such as self-avoidance and syncytial fusion, there may be other core proteins that are evolutionarily constrained. Conserved non-recognition proteins might also anchor other self-recognition proteins. By exploring external interactions in the multi-level context of the organism, population, and environment, we gain a better understanding of the different constraints on the evolution of self-recognition genes.

## Materials and Methods

### Bacterial strains and media

The strains and plasmids used in this study are described in Table 1. *P. mirabilis* strains were maintained on low swarm (LSW) agar (Belas et al., 1991). CM55 blood agar base agar (Oxoid, Basingstoke, England) was used for swarm-permissive nutrient plates. Overnight cultures of all strains were grown at 37°C in LB broth under aerobic conditions. For growth curve assays, cells were grown in minimal medium [M9 salts (3 g/L KH_2_PO_4_, 6.8 g/L Na_2_HPO_4_, 0.5 g/L NaCl, 1.0 g/L NH_4_Cl), 2 mM MgSO_4_, 0.1 mM CaCl_2_, 0.2% glucose] supplemented with 10mM L-serine (VWR, Beantown chemical, BT128350) when stated. Kanamycin (Corning, Corning, NY) was used at a concentration of 35 μg/ml for plasmid maintenance and was added to swarm and growth media when appropriate. Other antibiotics were used at the following concentrations for transforming plasmids into *P. mirabilis*: 15 μg/ml tetracycline (Amresco Biochemicals, Solon, OH), and 25 μg/ml streptomycin (Sigma-Aldrich, St. Louis, MO). Anhydrotetracycline (Sigma-Aldrich, St. Louis, MO) was used to induce gene expression from the Tet promoter at a concentration of 10 nM in the medium when stated.

**Table 1.** Strains used in this study.

| Strain | Name in this study | Description | Reference |
| --- | --- | --- | --- |
| *Proteus mirabilis* |  |  |  |
| BB2000 wild-type strain | BB2000 | wild-type *P. mirabilis*<br>BB2000 strain | Belas et al.,<br>1991 |
| BB2000 $\Delta ids$ carrying pldsBB$\Delta idsE$ | $\Delta ids$ pldsBB$\Delta idsE$ | BB2000 $\Delta ids$ carrying a vector expressing the *ids* locus from BB2000 but with *idsE* deleted (CCS06) | Saak & Gibbs, 2016 |
| BB2000 carrying empty vector | BB2000 empty vector | BB2000 carrying a plasmid without promoter-gene insert to confer antibiotic resistance | This study |
| BB2000 $\Delta idsE$ | $\Delta idsE$ | BB2000 with a chromosomal *idsE* deletion | Zepeda-Rivera et al.,<br>2018 |
| BB2000 $\Delta idsE$ carrying empty vector | $\Delta idsE$ empty vector | BB20000 $\Delta idsE$ carrying empty vector | This study |
| BB2000 $\Delta(idsE, sdaC)$ | ACH01 | BB20000 $\Delta idsE$ with a chromosomal *sdaC* (BB2000_0742) deletion | This study |
| BB2000 $\Delta(idsE, sdaC)$<br>carrying empty vector | ACH01 empty vector | ACH01 carrying empty<br>vector | This study |
| BB2000 $\Delta(idsE, sdaC)$<br>carrying pSdaC | ACH01 pSdaC | ACH01 carrying a plasmid<br>expressing *sdaC* from its<br>native promoter | This study |
| BB2000 $\Delta(sdaA,$<br>*sdaB)* | ACH04 | BB2000 with a<br>chromosomal deletion of<br>*sdaA* (BB2000_1697) and<br>*sdaB* (BB2000_0741) | This study |
| BB2000 $\Delta(sdaA,$<br>*sdaB)* carrying empty<br>vector | ACH04 empty vector | ACH04 carrying an empty<br>vector | This study |
| BB2000 $\Delta(sdaA,$<br>*sdaB-sdaC)* | ACH05 | ACH04 with a<br>chromosomal deletion of<br>*sdaC* | This study |
| BB2000 $\Delta(sdaA,$<br>*sdaB-sdaC)* carrying<br>empty vector | ACH05 empty vector | ACH05 carrying an empty<br>vector | This study |
| BB2000 $\Delta(sdaA,$<br>*sdaB, sstT)* | ACH04 $\Delta sstT$ | ACH04 with a<br>chromosomal deletion of<br>*sstT* (BB2000_0146) | This study |
| BB2000 $\Delta(sdaA, sdaB, yhaO)$ | ACH04 $\Delta yhaO$ | ACH04 with a chromosomal deletion of *yhaO* (BB2000_2747) | This study |
| BB2000 $\Delta(sdaA, sdaB-sdaC)$ carrying pSdaC | ACH05 pSdaC | ACH05 carrying pSdaC | This study |
| BB2000 $\Delta(idsE, sdaC)$ carrying pSdaC-I115A | ACH01 pSdaC-I115A | ACH01 carrying a modified pSdaC plasmid with I115A amino acid change | This study |
| BB2000 $\Delta(idsE, sdaC)$ carrying pSdaC-H325A | ACH01 pSdaC-H325A | ACH01 carrying a modified pSdaC plasmid with H325A amino acid change | This study |
| BB2000 $\Delta(idsE, sdaC)$ carrying pSdaC-V222W | ACH01 pSdaC-V222W | ACH01 carrying a modified pSdaC plasmid with V222W amino acid change | This study |
| BB2000 $\Delta(idsE, sdaC)$ carrying pSdaC-G328V | ACH01 pSdaC-G328V | ACH01 carrying a modified pSdaC plasmid with G328V amino acid change | This study |
| BB2000 $\Delta(idsE, sdaC)$<br>carrying pSdaC-G332R | ACH01 pSdaC-G332R | ACH01 carrying a<br>modified pSdaC plasmid<br>with G332R amino acid<br>change | This study |
| BB2000 $\Delta(sdaA, sdaB-sdaC)$ carrying<br>pSdaC-I115A | ACH05 pSdaC-I115A | ACH05 carrying a<br>modified pSdaC plasmid<br>with I115A amino acid<br>change | This study |
| BB2000 $\Delta(sdaA, sdaB-sdaC)$ carrying<br>pSdaC-H325A | ACH05 pSdaC-H325A | ACH05 carrying a<br>modified pSdaC plasmid<br>with H325A amino acid<br>change | This study |
| BB2000 $\Delta(sdaA, sdaB-sdaC)$ carrying<br>pSdaC-V222W | ACH05 pSdaC-V222W | ACH05 carrying a<br>modified pSdaC plasmid<br>with V222W amino acid<br>change | This study |
| BB2000 $\Delta(sdaA, sdaB-sdaC)$ carrying<br>pSdaC-G328V | ACH05 pSdaC-G328V | ACH05 carrying a<br>modified pSdaC plasmid<br>with G328V amino acid<br>change | This study |
| BB2000 $\Delta(sdaA, sdaB-sdaC)$ carrying pSdaC-G332R | ACH05 pSdaC-G332R | ACH05 carrying a modified pSdaC plasmid with G332R amino acid change | This study |
| BB2000 $\Delta(sdaA, sdaB-sdaC)$ carrying pTet-3xFLAG-SdaC | ACH05 pTet-SdaC-Pmir | ACH01 carrying a modified pSdaC plasmid with anhydrotetracycline inducible promoter and N-terminal 3xFLAG tag | This study |
| BB2000 $\Delta(sdaA, sdaB-sdaC)$ carrying pTet-SdaC-Ecoli | ACH05 pTet-SdaC-Ecoli | ACH01 carrying a plasmid expressing *E. coli* K12 MG1655 *sdaC* from an inducible anhydrotetracycline promoter | This study |
| BB2000 $\Delta(sdaA, sdaB-sdaC)$ carrying pTet-YhaO | ACH05 pTet-YhaO | ACH01 carrying a plasmid expressing *yhaO* from an inducible anhydrotetracycline promoter | This study |
| BB2000 $\Delta(sdaA, sdaB-sdaC)$ carrying pTet-YhaO-S234W | ACH05 pTet-YhaO-S234W | ACH01 carrying a plasmid expressing *yhaO* with S234W amino acid change from an inducible anhydrotetracycline promoter | This study |
| BB2000 $\Delta(idsE, sdaC)$ carrying pTet-3xFLAG-SdaC | ACH01 pTet-SdaC-Pmir | ACH01 carrying a modified pSdaC plasmid with anhydrotetracycline inducible promoter and N-terminal 3xFLAG tag | This study |
| BB2000 $\Delta(idsE, sdaC)$ carrying pTet-SdaC-Ecoli | ACH01 pTet-SdaC-Ecoli | ACH01 carrying a plasmid expressing *E. coli* K-12 MG1655 *sdaC* from an inducible anhydrotetracycline promoter | This study |
| BB2000 $\Delta(idsE, sdaC)$ carrying pTet-YhaO | ACH01 pTet-YhaO | ACH01 carrying a plasmid expressing *yhaO* from an inducible anhydrotetracycline promoter | This study |
| BB2000 $\Delta(idsE, sdaC)$<br>carrying pTet-3xFLAG-SdaC | ACH01 pTet-SdaC-Pmir-V222W | ACH01 carrying a<br>modified pSdaC plasmid<br>with anhydrotetracycline<br>inducible promoter and N-terminal 3xFLAG tag and<br>the V222W amino acid<br>change | This study |
| BB2000 $\Delta(idsE, sdaC)$<br>carrying pTet-YhaO-S234W | ACH01 pTet-YhaO-S234W | ACH01 carrying a plasmid<br>expressing *yhaO* with<br>S234W amino acid<br>change from an inducible<br>anhydrotetracycline<br>promoter | This study |
| *Escherichia coli* |  |  |  |
| One Shot Omnimax 2<br>T1R competent cells |  | Cloning strain for vectors | Thermo<br>Fisher<br>Scientific,<br>Waltham, MA |
| MFDpir |  | Mu-free mating strain for<br>*P. mirabilis* | Ferrières et<br>al., 2010 |
| SM10λpir | | Mating strain for moving<br>pKNG101 into *P. mirabilis* | Simon et al.,<br>1983 |

### Random mutagenesis of IdsE and spontaneous suppressor collection

Plasmids pIdsBB-IdsE-mut1 and pIdsBB-IdsE-mut2 were constructed by amplifying the *idsE* gene using oAC006 and oAC007 (Table S2) from the pIdsBB expression system containing a C-terminal GFPmut2 fusion (Gibbs et al., 2008) using error-prone PCR with the GeneMorph II Random Mutagenesis Kit (Agilent, Santa Clara, CA) and ligated back into the same pIdsBB expression vector using the restriction enzymes SacI and BamHI. Swarm-capable spontaneous mutants of the BB2000 Δ*ids* strains carrying pIdsBB-IdsE-mut1 or pIdsBB-IdsE-mut2 were isolated. Starting from frozen stocks, stable recovery of swarm expansion was verified. Plasmids were miniprepped and retransformed into the BB2000 Δ*ids* strain background to screen for plasmid-based suppressors, which were removed from further analysis. Boundary assays were performed to screen for defects in production or secretion of IdsD, and these suppressors were removed from further analysis. Seven of the remaining suppressor mutants were whole-genome sequenced (suppressors 1-7, Table S1).

The second set of swarm-capable spontaneous mutants were isolated from BB2000 Δ*idsE* (Zepeda-Rivera et al., 2018) carrying pTet-IdsE-mut3 and from ACH06 [BB2000 Δ(*sdaA*, *sdaB*, *idsE*)]. The *idsE*-mut3 sequence was amplified from pIdsBB-IdsE-T246A-S247A-T248A using oAC006 and oAC041 (Table S2) and ligated into the pTet vector (Zepeda-Rivera et al., 2018) using enzymes SacI and Bsu36I. Starting from frozen stocks, stable recovery of swarm expansion was verified. For suppressors derived from BB2000 Δ*idsE* pTet-IdsE-mut3, the promoter and gene were sequenced using Sanger sequencing (Genewiz, Inc., South Plainfield, NJ) to confirm causative mutations were chromosomal. Boundary assays were performed to screen for defects in production or secretion of IdsD, and these suppressor mutant strains were excluded from further analysis. Five of the remaining suppressor strains were whole-genome sequenced (suppressors 8-12, Table S1).

### Whole genome sequencing and variant calling

For the first round of sequencing (suppressors 1-7, Table S1), isolates were subjected to phenol-chloroform extractions to isolate genomic DNA (gDNA). gDNA was sheared using a Covaris S220 system (Covaris, Woburn, MA), and a library for whole-genome sequencing was prepared using the PrepX ILM DNA library kit (Takara Biosciences, Mountain View, CA) for the Apollo 324 next-generation sequencing (NGS) library prep system (Takara Biosciences). The library was sequenced as 75-bp paired-end reads using an Illumina NextSeq 500 system (Illumina, San Diego, CA). The Bauer Core Facility at Harvard University performed all genome sequencing.

For subsequent sequencing of the second set of mutants (suppressors 8-12, Table S1), gDNA was isolated as above, but library prep using KAPA HyperPrep kit (Roche, Wilmington, MA) was performed by the Bauer Core Facility. The library was sequenced as 150-bp paired-end reads using an Illumina NextSeq 500 system (Illumina, San Diego, CA) by the Bauer Core Facility. Breseq (Deatherage & Barrick, 2014) was used to perform variant calling of Illumina NextSeq reads against the BB2000 reference genome (GenBank accession no. CP004022).

### Plasmid construction

Restriction digestion was performed using restriction enzymes described (New England BioLabs, Ipswich, MA). Ligations were resolved in OneShot Omnimax2 T1R competent cells (Thermo Fisher Scientific, Waltham, MA) or SM10*λ*pir (Simon et al., 1983). The resultant plasmids were confirmed by Sanger sequencing (Genewiz, Inc., South Plainfield, NJ), and correct resultant plasmids were then transformed into *P. mirabilis* as described previously (Gibbs et al., 2008) using *E. coli* conjugative strain MFDpir (Ferrières et al., 2010). Table S2 contains the nucleotide sequences for listed primers, all of which contain the prefix “oAC.”

For pSdaC, the *sdaC* gene (BB2000_0742) including ~1kb upstream putative promoter region was amplified using oAC072 and oAC071 and ligated into the pTet vector (Zepeda-Rivera et al., 2018) using restriction enzymes NheI and PshAI to construct pSdaC. The empty vector is derived from pSdaC, containing no promoter or gene of interest but the rest of the backbone including kanamycin resistance and origin of replication is intact.

For pTet-SdaC-Pmir, the *sdaC* gene was amplified using oAC208 and oAC071 from pSdaC. Next, the sequence encoding the Tet promoter was amplified from pTet-FLAG-IdsE-mut3 with the addition of a 3XFLAG tag using oAC068, oAC196, oAC197, oAC198, and oAC207. The Tet promoter and *sdaC* gene were joined using overlap extension PCR with oAC068 and oAC071. The insert was ligated into the pTet vector using NheI and PshAI. For pTet-SdaC-Ecol, the *sdaC* gene was amplified from *E. coli* K-12 MG1655 using oAC177 and oAC178. Next, the Tet promoter sequence was amplified from the pTet vector using oAC68 and oAC185. The *sdaC* and Tet promoter sequences were joined using overlap extension PCR with oAC68 and oAC178. The insert was ligated into the pTet vector using NheI and Bsu36I. For pTet-YhaO, the *yhaO* gene (BB2000_2747) was amplified from *P. mirabilis* BB2000 using oAC187 and oAC188. Next, the Tet promoter sequence was amplified from the pTet vector using oAC68 and oAC186. The *yhaO* gene and Tet promoter sequence were joined using overlap extension PCR with oAC68 and oAC188. The insert was ligated into the pTet vector using NheI and Bsu36I.

SdaC variants were constructed by amplifying the *sdaC* gene from pSdaC using complementary primers containing the mutation along with oAC072 (SdaC native promoter) or oAC068 (Tet promoter) and oAC071 in overlap extension PCR. The *sdaC* gene containing the mutation was ligated back into the pSdaC vector using enzymes NheI and PshAI. The primers for each mutation were oAC264 and oAC265 for I115A, oAC268 and oAC269 for H325A, oAC293 and oAC294 for V222W, oAC118 and oAC119 for G332R, and oAC227 and oAC228 for G328V. The YhaO variant was constructed using complementary primers oAC314 and oAC315 containing the S324W mutation along with flanking primers oAC68 and oAC188 in overlap extension PCR. The insert was ligated into pTet-YhaO with enzymes NheI and Bsu36I.

### Strain construction

All chromosomal deletions were performed as described earlier using pKNG101-derived suicide vectors (Saak & Gibbs, 2016).

For strain ACH01 [BB20000 Δ(*idsE*, *sdaC*)], 500bp regions adjacent on either side to *sdaC* (BB2000_0742) with restriction sites were amplified using oAC046-049, and ligated into pKNG101 using restriction enzymes ApaI and SpeI. The resulting vector was mated into BB2000 Δ*idsE*. Matings were subjected to antibiotic selection on LSW agar (with 15 g/ml Tet and 25 g/ml Strep). Candidate strains were subjected to sucrose counterselection as described (Sturgill et al., 2002). Double recombinants were confirmed by PCR of the surrounding regions using oAC113 and oA114. ACH01 was confirmed by whole genome sequencing.

For strain ACH04 [BB2000 Δ(*sdaA*, *sdaB*)], the regions flanking the *sdaB* (BB2000_0741) gene were amplified using overlap extension PCR with oAC050-053 and ligated into pKNG101 using enzymes ApaI and SpeI. The resulting vector was mated into BB2000 and subjected to sucrose selection. Double recombinants were confirmed by colony PCR of the surrounding region using oAC113 and oAC114. The regions flanking the *sdaA* gene (BB2000_1697) were amplified using overlap extension PCR with oAC141-144 and ligated into pKNG101 using enzymes ApaI and SpeI. The resulting vector was mated into BB2000 Δ*sdaB* and subjected to sucrose selection. Double recombinants were confirmed by PCR of the surrounding region using oAC153 and oAC154.

For strain ACH05 [BB2000 Δ(*sdaA*, *sdaB*-*sdaC*)], the *sdaA* deletion vector used to construct ACH04 was mated into BB2000 and confirmed by PCR of the *sdaA* region using oAC153 and oAC154. The regions flanking the *sdaB-sdaC* genes were amplified using overlap extension PCR with oAC46, oAC53, oAC54, and oAC55 and ligated into pKNG101 using enzymes ApaI and SpeI. The resulting vector was mated into Δ*sdaA* and subjected to sucrose selection. Double recombinants were confirmed by PCR of the surrounding region using oAC113 and oAC114. Strain ACH05 was confirmed by whole genome sequencing.

For strain ACH04 Δ*sstT*, the regions flanking the *sstT* gene (BB2000_0146) were amplified using overlap extension PCR with oAC145-oAC148 and ligated into pKNG101 using enzymes ApaI and SpeI. The resulting vector was mated into ACH04 and subjected to sucrose selection. ACH04 Δ*sstT* was confirmed using PCR amplification of the surrounding region using oAC156 and oAC157. For strain ACH04 Δ*yhaO*, the regions flanking the *yhaO* gene (BB2000_2747) were amplified using overlap extension PCR with oAC161-oAC164 and ligated into pKNG101 using enzymes ApaI and SpeI. The resulting vector was mated into ACH04 and subjected to sucrose selection. ACH04 Δ*yhaO* was confirmed using PCR amplification of the surrounding region using oAC179 and oAC180.

### Growth curve

Overnight cultures were normalized to an optical density at 600 nm (OD600) of 0.1 in minimal medium [M9 salts (3 g/L KH_2_PO_4_, 6.8 g/L Na_2_HPO_4_, 0.5 g/L NaCl, 1.0 g/L NH_4_Cl), 2 mM MgSO_4_, 0.1 mM CaCl_2_, 0.2% glucose] supplemented with 10mM L-serine (VWR, Beantown chemical, BT128350) when stated. Both were supplemented with kanamycin for plasmid maintenance when appropriate. Normalized cultures were grown overnight at 37°C, with periodic shaking, in a Tecan Infinite 200 PRO microplate reader (Tecan, Männedorf, Switzerland).

### Swarm expansion

Overnight cultures were normalized to an OD600 of 0.1, and swarm-permissive nutrient plates supplemented with kanamycin were inoculated with 2 μl of normalized culture in the center. Plates were incubated at room temperature for two days, and the radii of actively migrating swarms starting from the edge of the inoculum were measured using Fiji (ImageJ) (Schindelin et al., 2012).

### LC-MS of L-serine in *P. mirabilis* cells

*P. mirabilis* cells were harvested by centrifugation from three swarm-permissive plates after incubation at 37°C for 16 to 20 hours. Cells were sequentially resuspended and centrifuged in 1ml of 100% LB, 80% LB, 60% LB, 40% LB, 20% LB, water (LC-MS grade water from Thermo Fisher Scientific, Waltham, MA). Cell pellets were flash frozen in liquid nitrogen and stored at −80°C. Pellets were resuspended in 1 ml cold LC-MS grade methanol (Sigma-Aldrich, St. Louis, MO) and lysed by vortexing for 10 minutes with cell disruptor beads (0.1-mm diameter; Electron Microscopy Sciences, Hatfield, PA). Lysate was transferred to an 8ml glass vial with an additional 1ml cold methanol rinse of the lysis tube. 4 mL of cold LC-MS grade chloroform (Sigma-Aldrich, St. Louis, MO) was added and samples were vortexed for 1 minute. 2 mL of LC-MS grade water containing 0.1 μL of MSK-A2-1.2 stable isotope-labeled amino acid standards (Cambridge Isotope Laboratories, Tewksbury, MA) was added before vortexing for 1 minute. After centrifuging for 10min at 3000rpm, 3.5 ml of the upper aqueous phase was transferred to a new glass vial and stored at −80°C. After removing the organic chloroform phase, the remaining interphase was dried completely before resuspending in Tris-buffered saline (50mM Tris-Cl, 150mM NaCl, pH 7.6) for protein quantification using a Bradford Assay (BioRad). Aqueous phase was evaporated under nitrogen flow and used for LC-MS, which was performed by the Small Molecule Mass Spectrometry Core Facility at Harvard University.

### Bioinformatics

We used I-TASSER (Roy et al., 2010; Yang et al., 2015; Y. Zhang, 2008) to predict the protein structure of SdaC-Pmir. Figures of structural models were made using PyMOL (The PyMOL Molecular Graphics System, Version 2.0 Schrödinger, LLC). Amino acid sequences in Figure 3 were aligned using Clustal Omega (Madeira et al., 2019; Sievers et al., 2011). Sequence alignments were visualized in Jalview (Waterhouse et al., 2009).

## Acknowledgments

We thank Charles Vidoudez and the Small Molecule Mass Spectrometry Core Facility at Harvard University for LC-MS serine concentration analysis as well as the Bauer Core Facility at Harvard University for assisting with whole-genome sequencing. Shamayeeta Ray provided much-appreciated technical advice on the design of SdaC variants. We thank members of the Gibbs, Cavanaugh, and Gaudet groups for insightful discussions.

## Author contributions

A.C. -- conceptualization, investigation, formal analysis, writing - original draft, reviewing and editing. K.A.G. -- conceptualization, formal analysis, supervision, project administration, writing - original draft, reviewing and editing.

## Data availability

Strains and primary data are available upon request.

## References

Armbruster, C. E., Mobley, H. L. T., & Pearson, M. M. (2018). Pathogenesis of *Proteus mirabilis* Infection. EcoSal Plus, 8(1), 10.1128/ecosalplus.ESP-0009-2017. https://doi.org/10.1128/ecosalplus.ESP-0009-2017

Asfahl, K. L., & Schuster, M. (2017). Social interactions in bacterial cell–cell signaling. FEMS Microbiology Reviews, 41(1), 92–107.https://doi.org/10.1093/femsre/fuw038

Baba, T., Ara, T., Hasegawa, M., Takai, Y., Okumura, Y., Baba, M., Datsenko, K. A., Tomita, M., Wanner, B. L., & Mori, H. (2006). Construction of *Escherichia coli* K-12 in-frame, single-gene knockout mutants: The Keio collection. Molecular Systems Biology, 2(1), 2006.0008.

Barroso-Batista, J., Pedro, M. F., Sales-Dias, J., Pinto, C. J. G., Thompson, J. A., Pereira, H., Demengeot, J., Gordo, I., & Xavier, K. B. (2020). Specific Eco-evolutionary Contexts in the Mouse Gut Reveal *Escherichia coli* Metabolic Versatility. Current Biology, 30(6), 1049–1062.e7. https://doi.org/10.1016/j.cub.2020.01.050

Belas, R., Erskine, D., & Flaherty, D. (1991). Transposon mutagenesis in *Proteus mirabilis*. J Bacteriol, 173(19), 6289–6293.

Bozzi, A. T., Zimanyi, C. M., Nicoludis, J. M., Lee, B. K., Zhang, C. H., & Gaudet, R. (2019). Structures in multiple conformations reveal distinct transition metal and proton pathways in an Nramp transporter. Elife, 8, e41124.

Brauer, A. L., White, A. N., Learman, B. S., Johnson, A. O., & Armbruster, C. E. (2019). D-Serine degradation by *Proteus mirabilis* contributes to fitness during single-species and polymicrobial catheter-associated urinary tract infection. MSphere, 4(1).

Cardarelli, L., Saak, C., & Gibbs, K. A. (2015). Two Proteins Form a Heteromeric Bacterial Self-Recognition Complex in Which Variable Subdomains Determine Allele-Restricted Binding. MBio, 6(3), e00251–15.

Chatterjee, S., & Rothenberg, E. (2012). Interaction of Bacteriophage l with Its *E. coli* Receptor, LamB. Viruses, 4(11), 3162–3178. https://doi.org/10.3390/v4113162

Connolly, J. P., Gabrielsen, M., Goldstone, R. J., Grinter, R., Wang, D., Cogdell, R. J., Walker, D., Smith, D. G., & Roe, A. J. (2016). A Highly Conserved Bacterial D-Serine Uptake System Links Host Metabolism and Virulence. PLoS Pathog, 12(1), e1005359. https://doi.org/10.1371/journal.ppat.1005359

de Jonge, P. A., Nobrega, F. L., Brouns, S. J., & Dutilh, B. E. (2019). Molecular and evolutionary determinants of bacteriophage host range. Trends in Microbiology, 27(1), 51–63.

de Oliveira, J. L., Morales, A. C., Stewart, B., Gruenheit, N., Engelmoer, J., Brown, S. B., de Brito, R. A., Hurst, L. D., Urrutia, A. O., & Thompson, C. R. (2019). Conditional expression explains molecular evolution of social genes in a microbe. Nature Communications, 10(1), 1–12.

Deatherage, D. E., & Barrick, J. E. (2014). Identification of mutations in laboratory-evolved microbes from next-generation sequencing data using breseq. In Engineering and analyzing multicellular systems (pp. 165–188). Springer.

Feldgarden, M., & Riley, M. A. (1999). The phenotypic and fitness effects of colicin resistance in *Escherichia coli* K-12. Evolution, 53(4), 1019–1027.

Ferrières, L., Hémery, G., Nham, T., Guérout, A.-M., Mazel, D., Beloin, C., & Ghigo, J.-M. (2010). Silent mischief: Bacteriophage Mu insertions contaminate products of *Escherichia coli* random mutagenesis performed using suicidal transposon delivery plasmids mobilized by broad-host-range RP4 conjugative machinery. Journal of Bacteriology, 192(24), 6418–6427.

Fischer, M. S., & Glass, N. L. (2019). Communicate and fuse: How filamentous fungi establish and maintain an interconnected mycelial network. Frontiers in Microbiology, 10, 619.

Gérard, F., Pradel, N., & Wu, L.-F. (2005). Bactericidal activity of colicin V is mediated by an inner membrane protein, SdaC, of *Escherichia coli*. Journal of Bacteriology, 187(6), 1945–1950.

Gibbs, K. A., Urbanowski, M. L., & Greenberg, E. P. (2008). Genetic determinants of self identity and social recognition in bacteria. Science, 321(5886), 256–259. https://doi.org/10.1126/science.1160033

Gonçalves, A. P., Heller, J., Rico-Ramírez, A. M., Daskalov, A., Rosenfield, G., & Glass, N. L. (2020). Conflict, Competition, and Cooperation Regulate Social Interactions in Filamentous Fungi. Annual Review of Microbiology, 74, 693–712.

Gruenheit, N., Parkinson, K., Stewart, B., Howie, J. A., Wolf, J. B., & Thompson, C. R. (2017). A polychromatic ‘greenbeard’ locus determines patterns of cooperation in a social amoeba. Nature Communications, 8(1), 1–9.

Hama, H., Kayahara, T., Tsuda, M., & Tsuchiya, T. (1991). Inhibition of homoserine dehydrogenase I by L-serine in *Escherichia coli*. The Journal of Biochemistry, 109(4), 604–608.

Hama, H., Shimamoto, T., Tsuda, M., & Tsuchiya, T. (1988). Characterization of a novel L-serine transport system in *Escherichia coli*. Journal of Bacteriology, 170(5), 2236–2239.

Hama, H., Sumita, Y., Kakutani, Y., Tsuda, M., & Tsuchiya, T. (1990). Target of serine inhibition in *Escherichia coli*. Biochemical and Biophysical Research Communications, 168(3), 1211–1216.

Hampton, H. G., Watson, B. N., & Fineran, P. C. (2020). The arms race between bacteria and their phage foes. Nature, 577(7790), 327–336.

Hirose, S., Chen, G., Kuspa, A., & Shaulsky, G. (2017). The polymorphic proteins TgrB1 and TgrC1 function as a ligand–receptor pair in *Dictyostelium* allorecognition. Journal of Cell Science, 130(23), 4002–4012.

Inglis, R. F., Scanlan, P., & Buckling, A. (2016). Iron availability shapes the evolution of bacteriocin resistance in *Pseudomonas aeruginosa*. The ISME Journal, 10(8), 2060.

Kearns, D. B. (2010). A field guide to bacterial swarming motility. Nature Reviews Microbiology, 8(9), 634–644.

Kitamoto, S., Alteri, C. J., Rodrigues, M., Nagao-Kitamoto, H., Sugihara, K., Himpsl, S. D., Bazzi, M., Miyoshi, M., Nishioka, T., & Hayashi, A. (2019). Dietary l-serine confers a competitive fitness advantage to *Enterobacteriaceae* in the inflamed gut. Nature Microbiology, 1–10.

Kleanthous, C. (2010). Swimming against the tide: Progress and challenges in our understanding of colicin translocation. Nature Reviews Microbiology, 8(12), 843–848.

Kostadinov, D., & Sanes, J. R. (2015). Protocadherin-dependent dendritic self-avoidance regulates neural connectivity and circuit function. Elife, 4, e08964.

Krishnamurthy, H., & Gouaux, E. (2012). X-ray structures of LeuT in substrate-free outward-open and apo inward-open states. Nature, 481(7382), 469.

Kundert, P., & Shaulsky, G. (2019). Cellular allorecognition and its roles in *Dictyostelium* development and social evolution. The International Journal of Developmental Biology, 63(8-9–10), 383.

Lefebvre, J. L., Kostadinov, D., Chen, W. V., Maniatis, T., & Sanes, J. R. (2012). Protocadherins mediate dendritic self-avoidance in the mammalian nervous system. Nature, 488(7412), 517–521.

Likhacheva, N. A., Samsonov, V. V., Samsonov, V. V., & Sineoky, S. P. (1996). Genetic control of the resistance to phage C1 of *Escherichia coli* K-12. J Bacteriol, 178(17), 5309–5315.

Little, K., Austerman, J., Zheng, J., & Gibbs, K. A. (2019). Cell shape and population migration are distinct steps of Proteus mirabilis swarming that are decoupled on high-percentage agar. Journal of Bacteriology, 201(11), e00726–18.

Madeira, F., Park, Y. M., Lee, J., Buso, N., Gur, T., Madhusoodanan, N., Basutkar, P., Tivey, A. R., Potter, S. C., & Finn, R. D. (2019). The EMBL-EBI search and sequence analysis tools APIs in 2019. Nucleic Acids Research, 47(W1), W636–W641.

Mangalea, M. R., & Duerkop, B. A. (2020). Fitness trade-offs resulting from bacteriophage resistance potentiate synergistic antibacterial strategies. Infection and Immunity.

Meaden, S., Paszkiewicz, K., & Koskella, B. (2015). The cost of phage resistance in a plant pathogenic bacterium is context-dependent. Evolution, 69(5), 1321–1328.

Molumby, M. J., Keeler, A. B., & Weiner, J. A. (2016). Homophilic protocadherin cell-cell interactions promote dendrite complexity. Cell Reports, 15(5), 1037–1050.

Nicoludis, J. M., Green, A. G., Walujkar, S., May, E. J., Sotomayor, M., Marks, D. S., & Gaudet, R. (2019). Interaction specificity of clustered protocadherins inferred from sequence covariation and structural analysis. Proceedings of the National Academy of Sciences, 116(36), 17825–17830.

Nicoludis, J. M., Lau, S.-Y., Schärfe, C. P., Marks, D. S., Weihofen, W. A., & Gaudet, R. (2015). Structure and sequence analyses of clustered protocadherins reveal antiparallel interactions that mediate homophilic specificity. Structure, 23(11), 2087–2098.

Nicoludis, J. M., Vogt, B. E., Green, A. G., Schärfe, C. P., Marks, D. S., & Gaudet, R. (2016). Antiparallel protocadherin homodimers use distinct affinity-and specificity-mediating regions in cadherin repeats 1–4. Elife, 5, e18449.

Noonan, J. P., Li, J., Nguyen, L., Caoile, C., Dickson, M., Grimwood, J., Schmutz, J., Feldman, M. W., & Myers, R. M. (2003). Extensive linkage disequilibrium, a common 16.7-kilobase deletion, and evidence of balancing selection in the human protocadherin α cluster. The American Journal of Human Genetics, 72(3), 621–635.

Oberai, A., Joh, N. H., Pettit, F. K., & Bowie, J. U. (2009). Structural imperatives impose diverse evolutionary constraints on helical membrane proteins. Proceedings of the National Academy of Sciences, 106(42), 17747–17750.

Ogawa, W., Kim, Y. M., Mizushima, T., & Tsuchiya, T. (1998). Cloning and expression of the gene for the Na+-coupled serine transporter from *Escherichia coli* and characteristics of the transporter. J Bacteriol, 180(24), 6749–6752.

Palmer, A. G., Ali, M., Yang, S., Parchami, N., Bento, T., Mazzella, A., Oni, M., Riley, M. C., Schneider, K., & Massa, N. (2016). Kin recognition is a nutrient-dependent inducible phenomenon. Plant Signaling & Behavior, 11(9), e1224045.

Pathak, D. T., Wei, X., Dey, A., & Wall, D. (2013). Molecular recognition by a polymorphic cell surface receptor governs cooperative behaviors in bacteria. PLoS Genet, 9(11), e1003891.

Roy, A., Kucukural, A., & Zhang, Y. (2010). I-TASSER: a unified platform for automated protein structure and function prediction. Nature Protocols, 5(4), 725.

Ruhe, Z. C., Nguyen, J. Y., Xiong, J., Koskiniemi, S., Beck, C. M., Perkins, B. R., Low, D. A., & Hayes, C. S. (2017). CdiA effectors use modular receptor-binding domains to recognize target bacteria. MBio, 8(2).

Ruhe, Z. C., Wallace, A. B., Low, D. A., & Hayes, C. S. (2013). Receptor polymorphism restricts contact-dependent growth inhibition to members of the same species. MBio, 4(4), e00480–13.

Saak, C. C., & Gibbs, K. A. (2016). The Self-Identity Protein IdsD Is Communicated between Cells in Swarming *Proteus mirabilis* Colonies. Journal of Bacteriology, 198(24), 3278–3286.

Schaffer, J. N., & Pearson, M. M. (2015). *Proteus mirabilis* and Urinary Tract Infections. Microbiol Spectr, 3(5). https://doi.org/10.1128/microbiolspec.UTI-0017-2013

Schindelin, J., Arganda-Carreras, I., Frise, E., Kaynig, V., Longair, M., Pietzsch, T., Preibisch, S., Rueden, C., Saalfeld, S., & Schmid, B. (2012). Fiji: An open-source platform for biological-image analysis. Nature Methods, 9(7), 676–682.

Shao, Z., Lin, R. T., & Newman, E. B. (1994). Sequencing and characterization of the *sdaC* gene and identification of the *sdaCB* operon in *Escherichia coli* K12. Eur J Biochem, 222(3), 901–907.

Shao, Z., & Newman, E. B. (1993). Sequencing and characterization of the *sdaB* gene from *Escherichia coli* K-12. European Journal of Biochemistry, 212(3), 777–784.

Sharir-Ivry, A., & Xia, Y. (2017). The impact of native state switching on protein sequence evolution. Molecular Biology and Evolution, 34(6), 1378–1390.

Sievers, F., Wilm, A., Dineen, D., Gibson, T. J., Karplus, K., Li, W., Lopez, R., McWilliam, H., Remmert, M., & Söding, J. (2011). Fast, scalable generation of high-quality protein multiple sequence alignments using Clustal Omega. Molecular Systems Biology, 7(1), 539.

Simon, R., Priefer, U., & Pühler, A. (1983). A broad host range mobilization system for in vivo genetic engineering: Transposon mutagenesis in gram negative bacteria. Bio/Technology, 1(9), 784–791.

Strassmann, J. E., Gilbert, O. M., & Queller, D. C. (2011). Kin discrimination and cooperation in microbes. Annu Rev Microbiol, 65, 349–367. https://doi.org/10.1146/annurev.micro.112408.134109

Sturgill, G. M., Siddiqui, S., Ding, X., Pecora, N. D., & Rather, P. N. (2002). Isolation of *lacZ* fusions to *Proteus mirabilis* genes regulated by intercellular signaling: Potential role for the sugar phosphotransferase (Pts) system in regulation. FEMS Microbiology Letters, 217(1), 43–50.

Su, H. S., Lang, B. F., & Newman, E. B. (1989). L-serine degradation in *Escherichia coli* K-12: Cloning and sequencing of the *sdaA* gene. Journal of Bacteriology, 171(9), 5095–5102.

Tipping, M. J., & Gibbs, K. A. (2019). Peer pressure from a *Proteus mirabilis* self-recognition system controls participation in cooperative swarm motility. PLoS Pathogens, 15(7), e1007885.

Velayudhan, J., Jones, M. A., Barrow, P. A., & Kelly, D. J. (2004). L-serine catabolism via an oxygen-labile L-serine dehydratase is essential for colonization of the avian gut by *Campylobacter jejuni*. Infection and Immunity, 72(1), 260–268.

Wall, D. (2014). Molecular recognition in myxobacterial outer membrane exchange: Functional, social and evolutionary implications. Molecular Microbiology, 91(2), 209–220.

Waterhouse, A. M., Procter, J. B., Martin, D. M., Clamp, M., & Barton, G. J. (2009). Jalview Version 2—A multiple sequence alignment editor and analysis workbench. Bioinformatics, 25(9), 1189–1191.

Wenren, L. M., Sullivan, N. L., Cardarelli, L., Septer, A. N., & Gibbs, K. A. (2013). Two independent pathways for self-recognition in *Proteus mirabilis* are linked by type VI-dependent export. MBio, 4(4), e00374–13. https://doi.org/10.1128/mBio.00374-13

West, S. A., Diggle, S. P., Buckling, A., Gardner, A., & Griffin, A. S. (2007). The social lives of microbes. Annu. Rev. Ecol. Evol. Syst., 38, 53–77.

Willett, J. L., Gucinski, G. C., Fatherree, J. P., Low, D. A., & Hayes, C. S. (2015). Contact-dependent growth inhibition toxins exploit multiple independent cell-entry pathways. Proceedings of the National Academy of Sciences, 112(36), 11341–11346.

Wu, Q. (2005). Comparative genomics and diversifying selection of the clustered vertebrate protocadherin genes. Genetics, 169(4), 2179–2188.

Yang, J., Yan, R., Roy, A., Xu, D., Poisson, J., & Zhang, Y. (2015). The I-TASSER Suite: Protein structure and function prediction. Nature Methods, 12(1), 7.

Zepeda-Rivera, M. A., Saak, C. C., & Gibbs, K. A. (2018). A proposed chaperone of the bacterial type VI secretion system functions to constrain a self-identity protein. J Bacteriol. https://doi.org/10.1128/JB.00688-17

Zhang, X., El-Hajj, Z. W., & Newman, E. (2010). Deficiency in L-serine deaminase interferes with one-carbon metabolism and cell wall synthesis in *Escherichia coli* K-12. J Bacteriol, 192(20), 5515–5525. https://doi.org/10.1128/JB.00748-10

Zhang, X., & Newman, E. (2008). Deficiency in l-serine deaminase results in abnormal growth and cell division of *Escherichia coli* K-12. Molecular Microbiology, 69(4), 870–881.

Zhang, Y. (2008). I-TASSER server for protein 3D structure prediction. BMC Bioinformatics, 9(1), 40.

Zhao, J., Gladieux, P., Hutchison, E., Bueche, J., Hall, C., Perraudeau, F., & Glass, N L. (2015). Identification of allorecognition loci in *Neurospora crassa* by genomics and evolutionary approaches. Molecular Biology and Evolution, 32(9), 2417–2432.

