## Supplementary information for "The conserved serine transporter SdaC moonlights to enable self recognition"

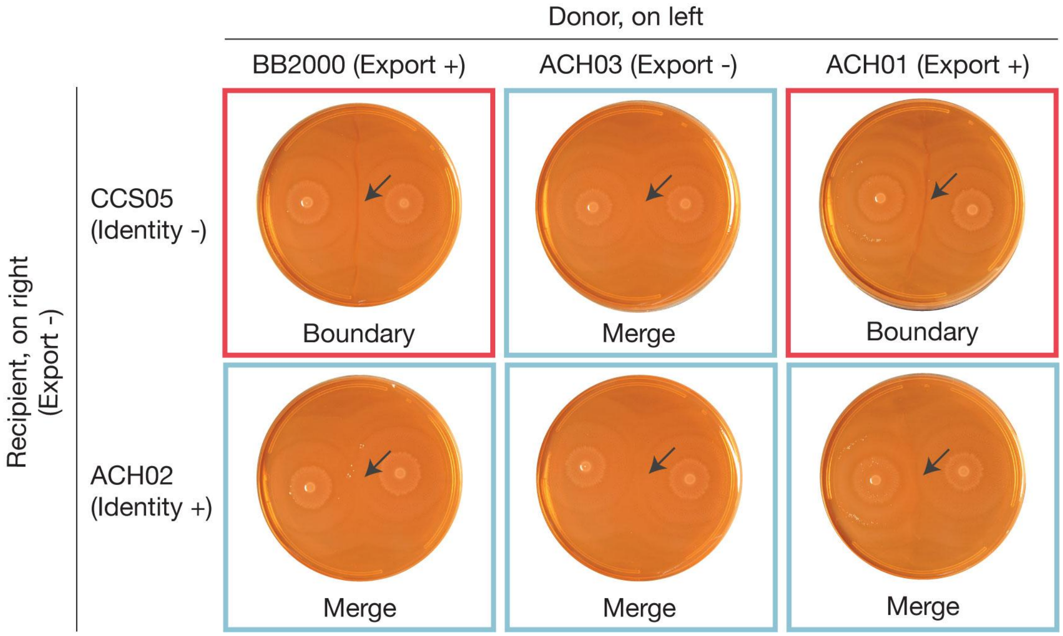

**Figure S1: The *sdaC* gene is not required for IdsD secretion between cells during swarming**

Boundary assay results shown for donor strains BB2000 (Export +), ACH03 (Export -), and ACH01 on the left side of the plate against recipient strains CCS05 (Identity -, Export -) and ACH02 (Identity +, Export -) on the right side of the plate. Intersection of strains is marked with a black arrow. Boundaries are outlined in a red box and merges are outlined in a blue box.

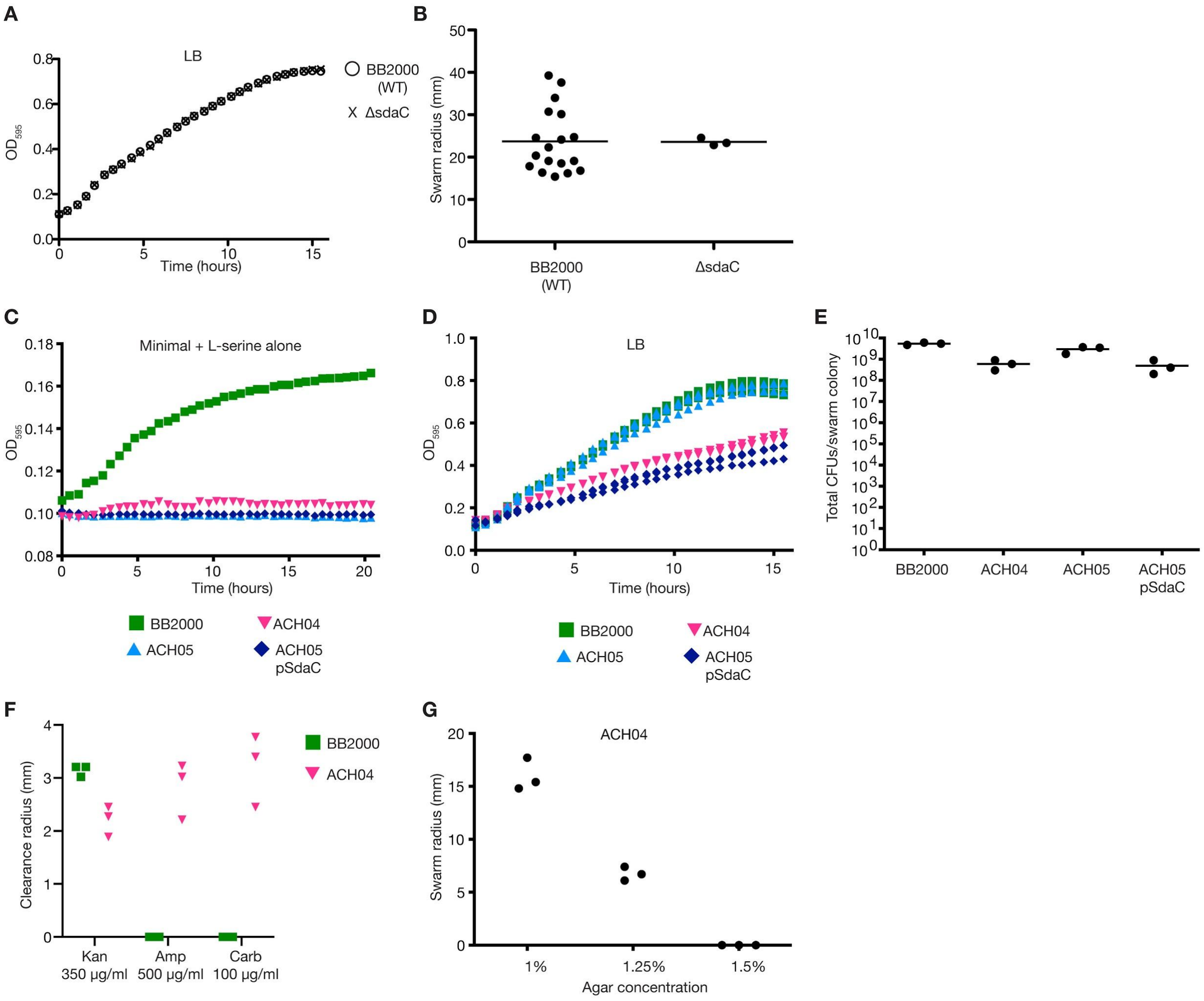

**Figure S2: SdaC activity in a high serine strain background reduces swarm expansion, slows growth, and destabilizes the cell wall**

**A)** Growth curve of BB2000 and BB2000 Δ*sdaC* in LB broth. **B)** Swarm radius measured from swarm assay of BB2000 Δ*sdaC* empty vector with BB2000 empty vector data copied from Figure 1C for comparison. **C)** Growth of BB2000 empty vector, ACH04 [BB2000 Δ(*sdaA*, *sdaB*)] empty vector, ACH05 [BB2000 Δ(*sdaA*, *sdaB*-*sdaC*)] empty vector, and ACH05 pSdaC in minimal medium without glucose and 10mM serine as the only carbon source. Averages shown for three biological replicates. **D)** Growth of BB2000 empty vector, ACH04 empty vector, ACH05 empty vector, and ACH05 pSdaC in LB for three biological replicates. **E)** CFUs per swarm colony for BB2000 empty vector, ACH04 empty vector, ACH05 empty vector, and ACH05 pSdaC for three biological replicates. **F)** Antibiotic clearance radius by discs soaked in Kan (350$\mu$g/ml), Amp (500$\mu$g/ml), or Carb (100$\mu$g/ml) for swarm colonies of strains BB2000 and ACH04. **G)** Swarm radius measured from swarm assay of ACH04 on LB medium containing 1%, 1.25%, or 1.5% agar.

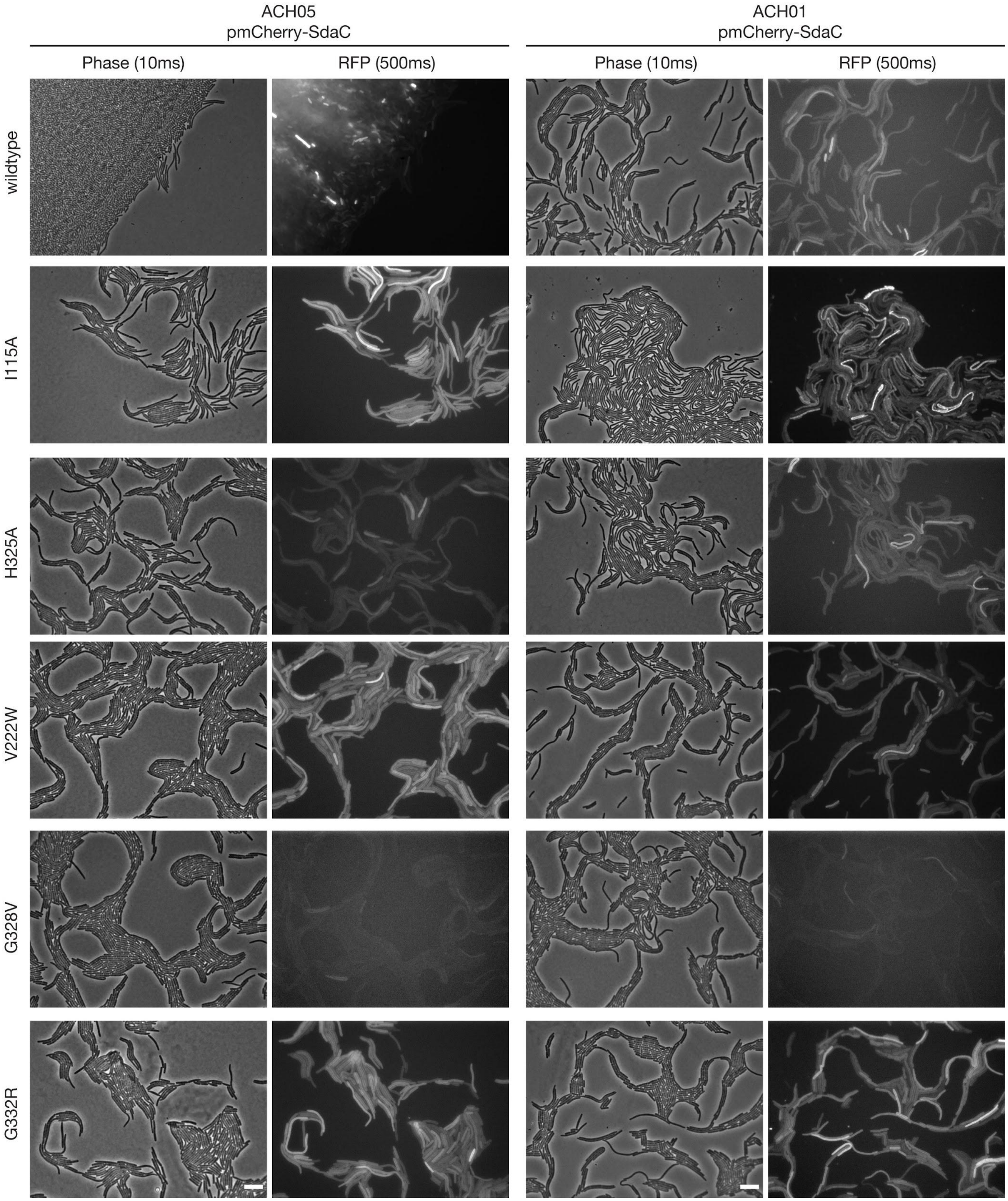

**Figure S3: Fluorescent fusion protein for wildtype SdaC and SdaC variants visible along the cell periphery**

Representative epifluorescence micrographs from phase and RFP channels of pmCherry-SdaC (wildtype or individual residue variants I115A, H325A, V222W, G328V, and G332R) expressed in ACH05 [BB2000 Δ(*sdaA*, *sdaB*-*sdaC*)] and ACH01 [BB2000 Δ(*idsE*, *sdaC*)]. Scale bar on the lower right of the last row of phase contrast images is 10$\mu$m.

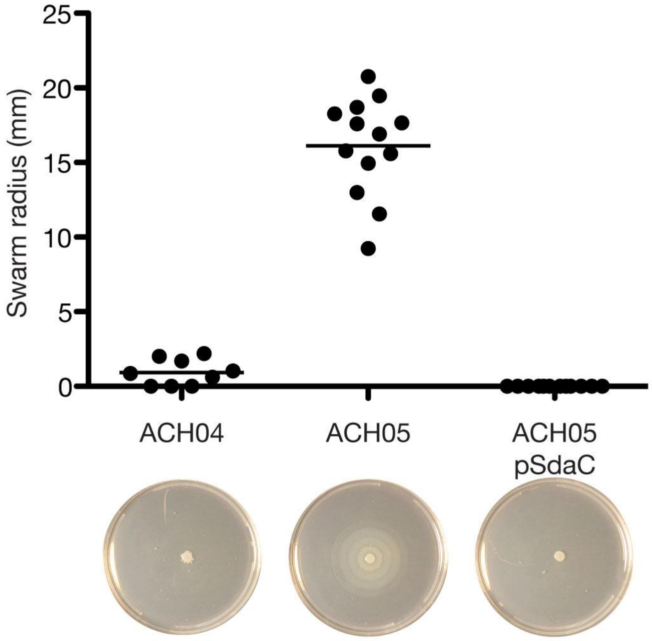

**Figure S4: SdaC activity restricts swarm expansion when *sdaA* and *sdaB* are deleted**
Representative swarm plate images for Figure 1G swarm assay of ACH04 [BB2000 Δ(*sdaA*, *sdaB*)] empty vector, ACH05 [BB2000 Δ(*sdaA*, *sdaB*-*sdaC*)] empty vector, and ACH05 pSdaC.

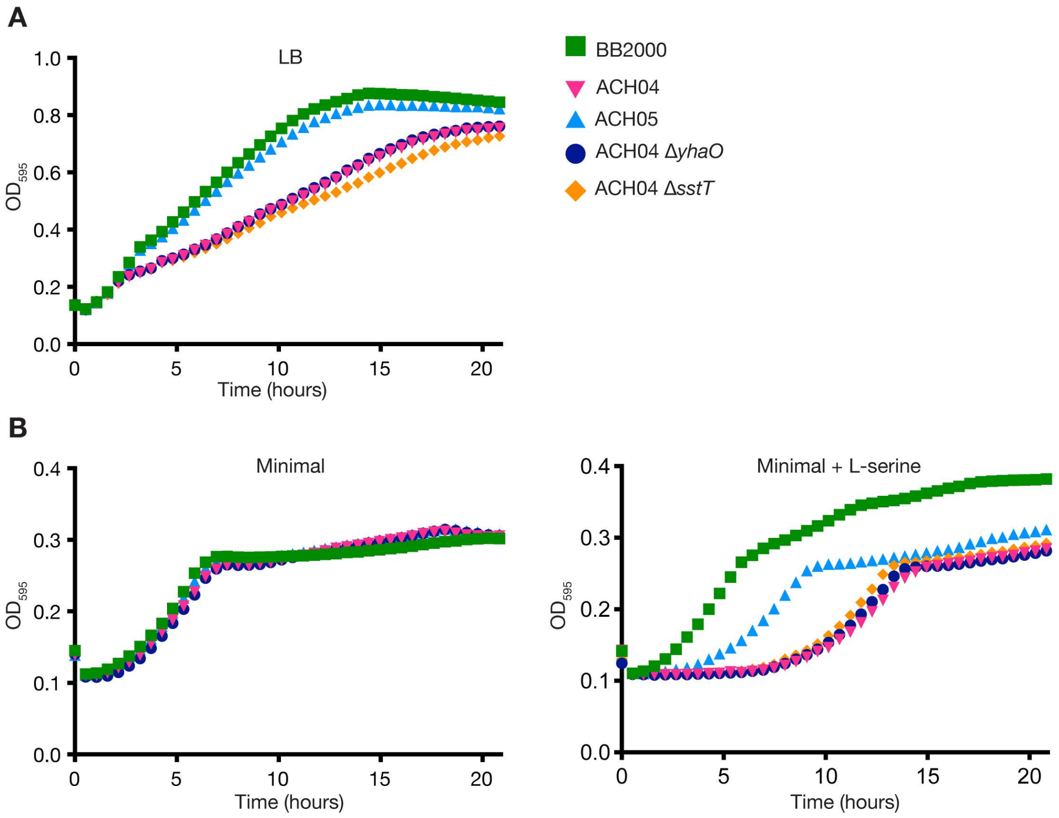

**Figure S5: Deletion of neither *sstT* nor *yhaO* in the ACH04 background rescues growth in LB or minimal medium compared to deletion of *sdaC***

**A)** Growth curve in LB of BB2000, ACH04 [BB2000 Δ(*sdaA*, *sdaB*)], ACH05 [BB2000 Δ(*sdaA*, *sdaB*-*sdaC*)], ACH04 Δ*sstT*, and ACH04 Δ*yhaO.* **B)** Growth curve in minimal medium (left) and minimal medium plus 10mM L-serine (right) of BB2000, ACH04, ACH05, ACH04 Δ*sstT*, and ACH04 Δ*yhaO.*

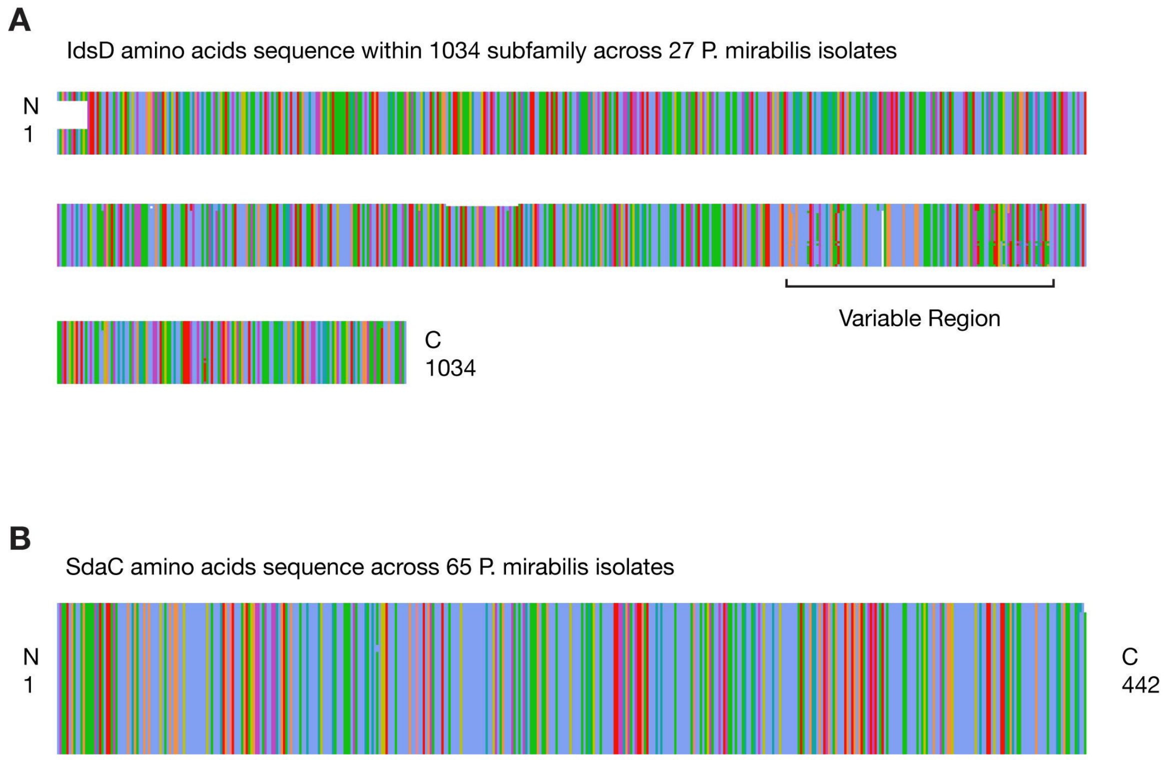

**Figure S6: IdsD is polymorphic across *P. mirabilis* isolates while SdaC is conserved throughout the protein sequence**
A) Alignment of 1034 subfamily IdsD amino acid sequences from 27 *P. mirabilis* isolate genomes available through NCBI. Alignment performed using MUSCLE and visualized with AlignmentViewer. B) Alignment of SdaC amino acid sequences from 65 *P. mirabilis* isolate genomes available through NCBI. Alignment performed using MUSCLE and visualized with AlignmentViewer.

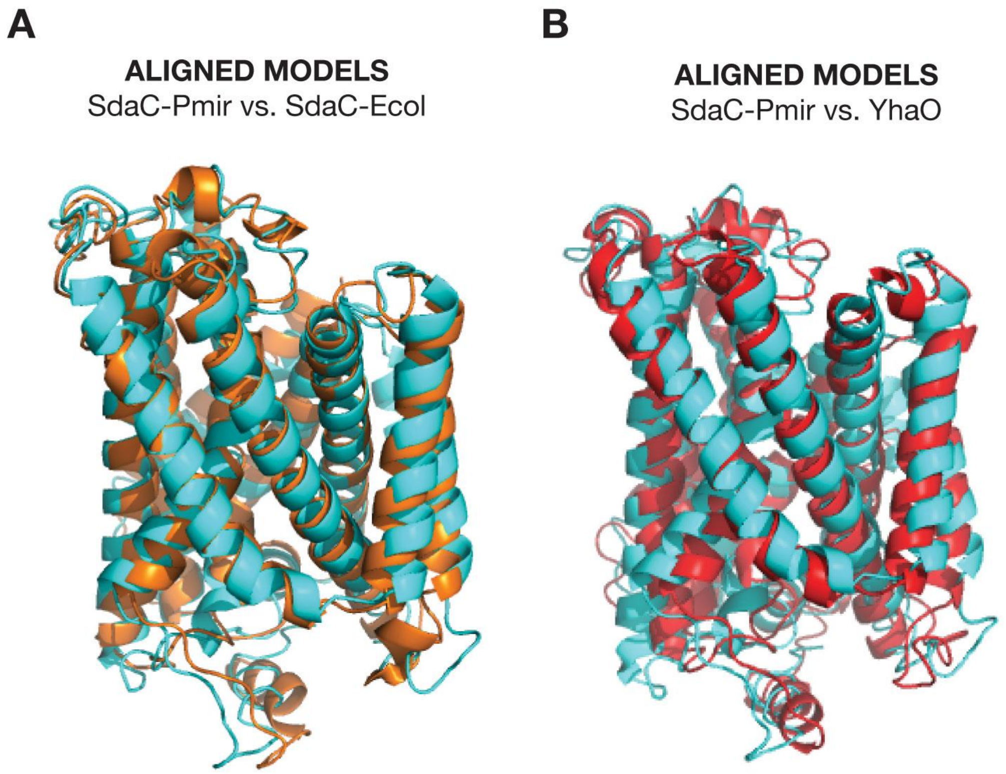

**Figure S7: Alignment of SdaC and YhaO I-TASSER models**
A) PyMol alignment of structural model of SdaC-Pmir from I-TASSER (blue) aligned to structural model of SdaC-Ecol from I-TASSER (orange) (C-score = -0.64, TM-score = 0.63±0.13, RMSD = 8.4±4.5Å). B) PyMol alignment of structural model of SdaC from I-TASSER (blue) aligned to structural model of YhaO from I-TASSER (red) (C-score = -0.55, TM-score = 0.64±0.13, RMSD = 8.3±4.5Å).

Table S1. Strains used in supplementary data.

| Strain | Name in this study | Description | Reference |
|---|---|---|---|
| *Proteus mirabilis* | | | |
| BB2000 | BB2000 | wild-type *P. mirabilis* BB2000 strain | Belas et al., 1991 |
| BB2000 Δ*ids* carrying pIdsBB-IdsE[K85N, L113V, N126K, I166N, N188Y, T248N, N260D]-GFP | BB2000 Δ*ids* pIdsBB-IdsE-mut1 | BB2000 Δ*ids* complemented with *ids* operon expressed from its native promoter where IdsE has a C-terminal GFPmut2 and contains mutations K85N, L113V, N126K, I166N, N188Y, T248N, N260D | This study |
| BB2000 Δ*ids* carrying pIdsBB-IdsE[L48Q, F64C, I208F, V230L]-GFP | BB2000 Δ*ids* pIdsBB-IdsE-mut2 | BB2000 Δ*ids* complemented with *ids* operon expressed from its native promoter where IdsE has a C-terminal GFPmut2 and contains mutations L48Q, F64C, I208F, V230L | This study |

| | | | |
|---|---|---|---|
| BB2000 Δ*idsE* carrying pTet-FLAG-IdsE[T246A, S247A, T248A] | BB2000 Δ*idsE* pTet-IdsE-mut3 | BB2000 Δ*idsE* carrying vector where IdsE contains an N-terminal FLAG tag and T246A, S247A, T248A mutations | This study |
| BB2000 carrying empty vector | BB2000 empty vector | BB2000 carrying a plasmid without promoter-gene insert to confer antibiotic resistance | This study |
| BB2000 Δ*ids* $tssB_{T95G}$ | CCS05 | BB2000 Δ*ids* $tssB_{T95G}$; deficient in T6SS-mediated transport | Saak & Gibbs, 2016 |
| BB2000 Δ*idsE* | Δ*idsE* | BB2000 with a chromosomal *idsE* deletion | Zepeda-Rivera et al., 2018 |
| BB2000 Δ(*idsE*, *sdaC*) | ACH01 | BB2000 Δ*idsE* with a chromosomal deletion of *sdaC* (BB2000_0742) | This study |
| BB2000 $tssB_{T95G}$ | ACH02 | BB2000 with a mutation in *tssB* (BB2000_0821) to inactivate type VI secretion | This study |

| | | | |
|---|---|---|---|
| BB2000 Δ*idsE* *tssB*$_{T95G}$ | ACH03 | BB2000 Δ*idsE* with a mutation in *tssB* (BB2000_0821) to inactivate type VI secretion | This study |
| BB2000 Δ(*sdaA, sdaB*) | ACH04 | BB2000 with chromosomal deletions of *sdaA* (BB2000_1697) and *sdaB* (BB2000_0741) | This study |
| BB2000 Δ(*sdaA, sdaB*) carrying empty vector | ACH04 empty vector | ACH04 carrying an empty vector | This study |
| BB2000 Δ(*sdaA, sdaB-sdaC*) | ACH05 | ACH04 with a chromosomal deletion of *sdaC* | This study |
| BB2000 Δ(*sdaA, sdaB-sdaC*) carrying empty vector | ACH05 empty vector | ACH05 carrying an empty vector | This study |
| BB2000 Δ(*sdaA, sdaB, sstT*) | ACH04 Δ*sstT* | ACH04 with a chromosomal deletion of *sstT* (BB2000_0146) | This study |

| | | | |
|---|---|---|---|
| BB2000 Δ(*sdaA, sdaB, yhaO*) | ACH04 Δ*yhaO* | ACH04 with a chromosomal deletion of *yhaO* (BB2000_2747) | This study |
| BB2000 Δ(*sdaA, sdaB-sdaC*) carrying pSdaC | ACH05 pSdaC | ACH05 carrying pSdaC | This study |
| BB2000 Δ(*sdaA, sdaB, idsE*) | AC06 | ACH04 with a chromosomal deletion of *idsE* | |
| BB2000 Δ*sdaC* | Δ*sdaC* | BB2000 with a chromosomal deletion of *sdaC* | This study |
| BB2000 Δ(*idsE*, *sdaC*) carrying pmCherry-SdaC | ACH01 pmCherry-SdaC | ACH01 carrying a modified pSdaC plasmid with an N-terminal mCherry fluorescent protein after the start codon | This study |
| BB2000 Δ(*idsE*, *sdaC*) carrying pmCherrySdaC-I115A | ACH01 pmCherrySdaC-I115A | ACH01 carrying pmCherry-SdaC with I115A amino acid change | This study |

| | | | |
|---|---|---|---|
| BB2000 Δ(*idsE*, *sdaC*) carrying pmCherrySdaC-H325A | ACH01 pmCherrySdaC-H325A | ACH01 carrying pmCherry-SdaC with H325A amino acid change | This study |
| BB2000 Δ(*idsE*, *sdaC*) carrying pmCherrySdaC-V222W | ACH01 pmCherrySdaC-V222W | ACH01 carrying pmCherry-SdaC with V222W amino acid change | This study |
| BB2000 Δ(*idsE*, *sdaC*) carrying pmCherrySdaC-G328V | ACH01 pmCherrySdaC-G328V | ACH01 carrying pmCherry-SdaC with G328V amino acid change | This study |
| BB2000 Δ(*idsE*, *sdaC*) carrying pmCherrySdaC-G332R | ACH01 pmCherrySdaC-G332R | ACH01 carrying pmCherry-SdaC with G332R amino acid change | This study |
| BB2000 Δ(*sdaA, sdaB-sdaC*) carrying pmCherry-SdaC | ACH05 pmCherry-SdaC | ACH05 carrying pmCherry-SdaC | This study |
| BB2000 Δ(*sdaA, sdaB-sdaC*) carrying pmCherrySdaC-I115A | ACH05 pmCherrySdaC-I115A | ACH05 carrying pmCherry-SdaC with I115A amino acid change | This study |

| | | | |
|---|---|---|---|
| BB2000 Δ(*sdaA, sdaB-sdaC*) carrying pmCherrySdaC-H325A | ACH05 pmCherrySdaC-H325A | ACH05 carrying pmCherry-SdaC with H325A amino acid change | This study |
| BB2000 Δ(*sdaA, sdaB-sdaC*) carrying pmCherrySdaC-V222W | ACH05 pmCherrySdaC-V222W | ACH05 carrying pmCherry-SdaC with V222W amino acid change | This study |
| BB2000 Δ(*sdaA, sdaB-sdaC*) carrying pmCherrySdaC-G328V | ACH05 pmCherrySdaC-G328V | ACH05 carrying pmCherry-SdaC with G328V amino acid change | This study |
| BB2000 Δ(*sdaA, sdaB-sdaC*) carrying pmCherrySdaC-G332R | ACH05 pmCherrySdaC-G332R | ACH05 carrying pmCherry-SdaC with G332R amino acid change | This study |

| Suppressor strain background | SdaC mutation | Description | Reference |
|---|---|---|---|
| BB2000 Δ*ids* pIdsBB-IdsE[K85N, L113V, N126K, I166N, N188Y, T248N, N260D]-GFP | L371* | Suppressor 1, results in non-functional SdaC | This study |

| | | | |
|---|---|---|---|
| BB2000 Δ*ids* pIdsBB-IdsE[K85N, L113V, N126K, I166N, N188Y, T248N, N260D]-GFP | Δ68bp (366-433) | Suppressor 2, results in non-functional SdaC | This study |
| BB2000 Δ*ids* pIdsBB-IdsE[K85N, L113V, N126K, I166N, N188Y, T248N, N260D]-GFP | E243* | Suppressor 3, results in non-functional SdaC | This study |
| BB2000 Δ*ids* pIdsBB-IdsE[L48Q, F64C, I208F, V230L]-GFP | Δ1bp (244) | Suppressor 4, results in non-functional SdaC | This study |
| BB2000 Δ*ids* pIdsBB-IdsE[L48Q, F64C, I208F, V230L]-GFP | G332R | Suppressor 5, results in full-length SdaC with single residue change | This study |
| BB2000 Δ*ids* pIdsBB-IdsE[L48Q, F64C, I208F, V230L]-GFP | Δ1bp (200) | Suppressor 6, results in non-functional SdaC | This study |
| BB2000 Δ*ids* pIdsBB-IdsE[L48Q, F64C, I208F, V230L]-GFP | Δ49bp (575-623) | Suppressor 7, results in non-functional SdaC | This study |
| BB2000 Δ(*sdaA, sdaB, idsE*) | +2bp (363) | Suppressor 8, results in non-functional SdaC | This study |

| | | | |
|---|---|---|---|
| BB2000 Δ(*sdaA, sdaB, idsE*) | G192* | Suppressor 9, results in non-functional SdaC | This study |
| BB2000 Δ(*sdaA, sdaB, idsE*) | Δ2194bp (all sdaC plus 751bp upstream and 123bp downstream of sdaB) | Suppressor 10, results in non-functional SdaC | This study |
| BB2000 Δ*idsE* pTet-FLAG-IdsE[T246A, S247A, T248A] | G328V | Suppressor 11, results in full-length SdaC with single residue change | This study |
| BB2000 Δ*idsE* pTet-FLAG-IdsE[T246A, S247A, T248A] | Δ1bp (725) | Suppressor 12, results in non-functional SdaC | This study |

Table S2. Primers used for plasmid and strain construction

| **Primer name** | **Primer sequence** |
|---|---|
| oAC006 | GGCCCATGCCTGAGCTCGATT |
| oAC007 | CAGCTGATCCGGATCCCGCA |
| oAC041 | ACTACCTCAGGGATACTACGCATGG |
| oAC046 | ATCGGGGCCCGCGAAAGTTAAAATAATGTTTTA |
| oAC047 | TAAACTTAAATACTCTCCGAAATAACGCGGT |
| oAC048 | AGAGTATTTAAGTTTATCGATGGCTACTTTC |
| oAC049 | ATTAACTAGTATAGTTCGCCATTATAGGCGCTG |
| oAC050 | ATTAGGGCCCATACATTGCACCTTTCATTGCCT |
| oAC051 | GTGTTAAAAATATTCGCCTCCCTAATTTAAAGGC |
| oAC052 | CGAATATTTTTAACACTTTGACATGATTGTTACCCA |
| oAC053 | ATTAACTAGTATAAGCACCAATTTACCCTGTTT |
| oAC054 | GTGTTAAAAATACTCTCCGAAATAACGCGGTAA |
| oAC055 | AGAGTATTTTTAACACTTTGACATGATTGTTAC |
| oAC068 | AAATGGCTAGCTTAAGACCCACTTTCACATTTAAG |
| oAC071 | CGCCGACACGGGTCACGCTG |
| oAC072 | ATGGGCAAATGGCTAGCTCCTATTAGGATAAGAAA |
| oAC113 | GTGATCACGGTAGTTTATCC |
| oAC114 | TGTAGAGCTACATGAGTGAT |
| oAC118 | TTTAGGGGCTCGTGAACGTTTCAATGGCTTAAT |
| oAC119 | ATTAAGCCATTGAAACGTTCACGAGCCCCTAAA |
| oAC139 | GCATACTCATACAAGGAGCTTA |

| | |
|---|---|
| oAC140 | GCTCTTGAGCTTCTGCATTT |
| oAC141 | AACCCGGGCCCATATCCCAGTATGGTTTCTGTGAT |
| oAC142 | ATAAAATCAGGCTAAGTCGAAAACGCTAATCACGT |
| oAC143 | ATTAGCGTTTTCGACTTAGCCTGATTTTATTTCTA |
| oAC144 | ACTATACTAGTATATAATGAGATTTATTTTAAGACAGGCTACTCC |
| oAC145 | AACCCGGGCCCATATTTCCAGTTTAGGATACGTTG |
| oAC146 | TACTACTTATACTCTAGTTATTCCATTACTTAAAATAATATTTAAAAAAGCAATT |
| oAC147 | AGTAATGGAATAACTAGAGTATAAGTAGTAAATAGATATAAAGAGTAGTAAGACA |
| oAC148 | ACTATACTAGTATATTTGTGTGAGCTTGATCAAACACCAA |
| oAC153 | TCTCATGCCGGACAAGTTGC |
| oAC154 | ACAATTGACCCCCAACCAAA |
| oAC156 | GGCTTCCTAAAACTGAGTCA |
| oAC157 | GCAGGGAACAGAATTAGCAC |
| oAC161 | AACCCGGGCCCATATTATATAATTAACACTCTTTT |
| oAC162 | AACCGCAGTTGACGAATTAGTACTCACTTTTTATATTGTAATTGT |
| oAC163 | AAAGTGAGTACTAATTCGTCAACTGCGGTTCGAGTTAGCTGTTAT |
| oAC164 | ACTATACTAGTATATTAGCAATACTGACAATGGCATAACCTGACT |

| | |
|---|---|
| oAC177 | ATTTCGGAGAGTATTATGGAAACGACTCAAACCAG |
| oAC178 | CTGGCCTCAGGGATATTAGCTGAACAGAGAGTAGA |
| oAC179 | CGGTTTACGCACTTCTTGTC |
| oAC180 | TCTTCAGCACAAACCGTCGC |
| oAC185 | TTGAGTCGTTTCCATAATACTCTCCGAAATTTTCTCTATC |
| oAC186 | ATTAGTACTCACTTTTTTCTCTATCACTGATAGGGAGTGGTAA AA |
| oAC187 | TCAGTGATAGAGAAAAAAGTGAGTACTAATATGGAAACGGCT TCC |
| oAC188 | CTGGCCTCAGGGATACTATGAGAATGCCAAGAACGGAGAAA CACA |
| oAC196 | ATCGTCTTTGTAGTCTTTATCGTCATCGTCTTTGTAGTCC |
| oAC197 | GTAGTCTTTATCGTCATCGTCTTTGTAGTCTTTATCGTCA |
| oAC198 | TTGTATCTTTATCGTCATCGTCTTTGTAGTCTTTATCGTC |
| oAC207 | CCCTGTGCGATAGAACCAGCTTTGGTTGTATCTTTATCGT |
| oAC208 | TACAACCAAAGCTGGTTCTATCGCACAGGGTAATA |
| oAC226 | CTGTGCGATAGAACCAGCTTTGGTTGTATCCTTGTACAGCTC GTCCATGC |
| oAC227 | TTAGGTCACTATTTAGTGGCTCGTGAAGGTTTC |
| oAC228 | GAAACCTTCACGAGCCACTAAATAGTGACCTAA |
| oAC264 | TTTGCTATTTATCCTGCCTTGTTGGTTTACAGT |
| oAC265 | ACTGTAAACCAACAAGGCAGGATAAATAGCAAA |
| oAC268 | AAATCTTTCTTAGGTGCCTATTTAGGGGCTCGT |

| | |
|---|---|
| oAC269 | ACGAGCCCCTAAATAGGCACCTAAGAAAGATTT |
| oAC293 | TGGTTAGCTATCCCTTGGATGGTGTTCTCTTTT |
| oAC294 | AAAAGAGAACACCATCCAAGGGATAGCTAACCA |
| oAC295 | CTCGCCCTTGCTCACCATAATACTCTCCGAAATAACGCGG |
| oAC296 | TCGGAGAGTATTATGGTGAGCAAGGGCGAGGAGGATAACA |
| oAC314 | CACTACCTTTTACATTAACTTGGATTTTGTTTATCCAAACATT |
| oAC315 | AATGTTTGGATAAACAAAATCCAAGTTAATGTAAAAGGTAGTG |

Materials and Methods:

**Bacterial strains and media**

The strains and plasmids used in the supplementary information are described in Table S1. *P. mirabilis* strains were maintained on low swarm (LSW) agar (Belas et al., 1991). CM55 blood agar base agar (Oxoid, Basingstoke, England) was used for swarm-permissive nutrient plates, except when LB agar was used as a substitute in Figure S2G to vary agar concentration. Overnight cultures of all strains were grown at 37°C in LB broth under aerobic conditions. For growth curve assays, cells were grown in LB or minimal medium [M9 salts (3 g/L $KH_2PO_4$, 6.8 g/L $Na_2HPO_4$, 0.5 g/L NaCl, 1.0 g/L $NH_4Cl$), 2 mM $MgSO_4$, 0.1 mM $CaCl_2$, 0.2% glucose] supplemented with 10mM L-serine (VWR, Beantown chemical, BT128350) when stated. Glucose was omitted from the minimal medium in Figure S2C. Kanamycin (Corning, Corning, NY) was used at a concentration of 35 µg/ml for plasmid maintenance and was added to swarm and growth media when appropriate. Other antibiotics were used as follows for transforming plasmids into *P. mirabilis*: 15 µg/ml tetracycline (Amresco Biochemicals, Solon, OH), and 25 µg/ml streptomycin (Sigma-Aldrich, St. Louis, MO).

**Plasmid construction**

Restriction digestion using restriction enzymes described (New England BioLabs, Ipswich, MA). Ligations were resolved in OneShot Omnimax2 T1R competent cells (Thermo Fisher Scientific, Waltham, MA) or SM10*λ*pir (Simon et al., 1983). The resultant plasmids were confirmed by Sanger sequencing (Genewiz, Inc., South Plainfield, NJ), and correct resultant plasmids were then transformed into *P. mirabilis* as described previously (Gibbs et al., 2008) using *E. coli* conjugative strain MFDpir (Ferrières et al., 2010). Table S2 contains the nucleotide sequences for listed primers, all of which contain the prefix "oAC."

For pmCherry-SdaC, the native promoter for the *sdaC* gene was amplified from pSdaC using oAC072 and oAC295. Next, the gene encoding mCherry was amplified using oAC296 and oAC226. SdaC was amplified from pSdaC using oAC208 and oAC071. PCR products were joined together using overlap extension PCR with oAC072 and oAC071. The full insert was ligated into the pSdaC backbone using NheI and PshAI. Point mutations were made in the pmCherry-SdaC as described for point mutations in pSdaC using the same primers.

**Strain construction**

All chromosomal deletions were performed as described earlier using pKNG101-derived suicide vectors (Saak & Gibbs, 2016).

For strains ACH02 (BB2000 $tssB_{T95G}$) and ACH03 (BB2000 $tssB_{T95G}$ Δ*idsE*), pCS34 (Saak & Gibbs, 2016) was mated into BB2000 to construct ACH02 and BB2000 Δ*idsE* to construct ACH03. Matings were subjected to antibiotic selection on LSW agar (with 15 g/ml Tet and 25 g/ml Strep). Candidate strains were subjected to sucrose counterselection as described (Sturgill et al., 2002). Double recombinants were confirmed by PCR of the SNP-containing region of *tssB* (BB2000_0821) using oAC139 and oAC140.

For strain BB2000 Δ*sdaC*, 500bp regions adjacent on either side to *sdaC* with restriction sites were amplified using oAC046-049, and ligated into pKNG101 using restriction enzymes ApaI and SpeI. The resulting vector was mated into BB2000. Matings were subjected to antibiotic selection on LSW agar (with 15 g/ml Tet and 25 g/ml Strep). Candidate strains were subjected to sucrose counterselection as described (Sturgill et al., 2002). For BB2000 Δ*sdaC*, double recombinants were confirmed by PCR using oAC113 and oA114.

For strain ACH06 [BB2000 Δ(*sdaA*, *sdaB*, *idsE)*], the regions flanking the *sdaB* gene were amplified using overlap extension PCR with oAC050-053 and ligated into pKNG101 using enzymes ApaI and SpeI. The resulting vector was mated into BB2000 Δ*idsE* and subjected to sucrose selection. Double recombinants were confirmed by colony PCR of the region using oAC113 and oAC114. The regions flanking the *sdaA* gene were amplified using overlap extension PCR with oAC141-144 and ligated into pKNG101 using enzymes ApaI and SpeI. The resulting vector was mated into BB2000 Δ(*idsE, sdaB*) and subjected to sucrose selection. Double recombinants were confirmed by PCR of the surrounding region using oAC153 and oAC154. ACH06 was confirmed by whole genome sequencing.

**Growth curve**

Overnight cultures were normalized to an an optical density at 600 nm (OD600) of 0.1 in LB medium or minimal medium [M9 salts (3 g/L $KH_2PO_4$, 6.8 g/L $Na_2HPO_4$, 0.5 g/L NaCl, 1.0 g/L $NH_4Cl$), 2 mM $MgSO_4$, 0.1 mM $CaCl_2$, 0.2% glucose] supplemented with

10mM L-serine (VWR, Beantown chemical, BT128350) when stated. Glucose was omitted from the minimal medium in Figure S2C. Medium was supplemented with kanamycin for plasmid maintenance when appropriate. Normalized cultures were grown overnight at 37°C, with periodic shaking, in a Tecan Infinite 200 PRO microplate reader (Tecan, Männedorf, Switzerland).

**Swarm expansion and boundary assay**

Overnight cultures were normalized to an OD600 of 0.1, and swarm-permissive nutrient plates supplemented with kanamycin were inoculated with 2 µl of normalized culture in the center. For Figure S2G, LB medium with various percentages of agar was substituted for CM55 (Oxoid, Basingstoke, England) as the swarm-permissive nutrient medium. Plates were incubated at room temperature for two days, and the radii of actively migrating swarms starting from the edge of the inoculum were measured using Fiji (ImageJ) (Schindelin et al., 2012). Colony-forming units (CFUs) per swarm colony were calculated by resuspending all of the cells from a single plate in LB at the end of the swarm assay experiment and plating dilutions onto LSW agar plates that were incubated overnight at 37°C for growth of single colonies. Boundary assays were performed as reported (Wenren et al., 2013).

**Microscopy**

One-millimeter-thick swarm-permissive agar pads supplemented with kanamycin were inoculated with overnight cultures and incubated overnight at room temperature. The agar pads were then incubated at 37°C in a modified humidity chamber. After five to six hours, the pads were imaged by phase-contrast microscopy (10 ms exposure) and fluorescence microscopy (RFP channel, 500 ms exposure) using a Leica DM5500B microscope system (Leica Microsystems, Buffalo Grove, IL) and a CoolSnap HQ2 cooled charge-coupled device camera (Photometrics, Tucson, AZ). MetaMorph (version 7.8.0.0; Molecular Devices, Sunnyvale, CA) was used for image acquisition.

**Antibiotic clearance assay**

50 µl of overnight culture was spread evenly across a swarm-permissive nutrient plate. 6-mm blank paper disks were soaked into antibiotics [Kan (350$\mu$g/ml), Amp (500$\mu$g/ml), or Carb (100$\mu$g/ml)] or water as a control before placing on plates. After incubating plates at 37°C overnight, the radius of clearance surrounding the disk was measured.

**Bioinformatics**

I-TASSER (Roy et al., 2010; Yang et al., 2015; Zhang, 2008) was used to predict the protein structure of SdaC-Pmir, SdaC-Ecol, and YhaO. Figures aligning structural models were made using PyMOL (The PyMOL Molecular Graphics System, Version 2.0 Schrödinger, LLC). Additional IdsD and SdaC sequences from *P. mirabilis* genomes were identified by using tblastn (NCBI) with the *P. mirabilis* amino acid sequence as the query. Sequences with less than 90% query coverage and IdsD sequences from the 1072 subfamily were excluded. Sequence alignments were generated using MUSCLE (Edgar, 2004; Madeira et al., 2019). AlignmentViewer (Reguant et al., 2020) was used to visualize MUSCLE alignments of IdsD and SdaC for Figure S6.

### References

Belas, R., Erskine, D., & Flaherty, D. (1991). Transposon mutagenesis in *Proteus mirabilis*. *J Bacteriol*, *173*(19), 6289–6293.

Edgar, R. C. (2004). MUSCLE: multiple sequence alignment with high accuracy and high throughput. *Nucleic Acids Research*, *32*(5), 1792–1797.

Ferrières, L., Hémery, G., Nham, T., Guérout, A.-M., Mazel, D., Beloin, C., & Ghigo, J.-M. (2010). Silent mischief: Bacteriophage Mu insertions contaminate products of *Escherichia coli* random mutagenesis performed using suicidal transposon delivery plasmids mobilized by broad-host-range RP4 conjugative machinery. *Journal of Bacteriology*, *192*(24), 6418–6427.

Gibbs, K. A., Urbanowski, M. L., & Greenberg, E. P. (2008). Genetic determinants of self identity and social recognition in bacteria. *Science*, *321*(5886), 256–259. https://doi.org/10.1126/science.1160033

Madeira, F., Park, Y. M., Lee, J., Buso, N., Gur, T., Madhusoodanan, N., Basutkar, P., Tivey, A. R., Potter, S. C., & Finn, R. D. (2019). The EMBL-EBI search and sequence analysis tools APIs in 2019. *Nucleic Acids Research*, *47*(W1), W636–W641.

Reguant, R., Antipin, Y., Sheridan, R., Dallago, C., Diamantoukos, D., Luna, A., Sander, C., & Gauthier, N. P. (2020). AlignmentViewer: Sequence Analysis of Large Protein Families. *F1000Research*, *9*(213), 213.

Roy, A., Kucukural, A., & Zhang, Y. (2010). I-TASSER: a unified platform for automated protein structure and function prediction. *Nature Protocols*, *5*(4), 725.

Saak, C. C., & Gibbs, K. A. (2016). The Self-Identity Protein IdsD Is Communicated between Cells in Swarming *Proteus mirabilis* Colonies. *Journal of Bacteriology*, *198*(24), 3278–3286.

Schindelin, J., Arganda-Carreras, I., Frise, E., Kaynig, V., Longair, M., Pietzsch, T., Preibisch, S., Rueden, C., Saalfeld, S., & Schmid, B. (2012). Fiji: An open-source platform for biological-image analysis. *Nature Methods*, *9*(7), 676–682.

Simon, R., Priefer, U., & Pühler, A. (1983). A broad host range mobilization system for in vivo genetic engineering: Transposon mutagenesis in gram negative bacteria. *Bio/Technology*, *1*(9), 784–791.

Sturgill, G. M., Siddiqui, S., Ding, X., Pecora, N. D., & Rather, P. N. (2002). Isolation of *lacZ* fusions to *Proteus mirabilis* genes regulated by intercellular signaling: Potential role for the sugar phosphotransferase (Pts) system in regulation. *FEMS Microbiology Letters*, *217*(1), 43–50.

Wenren, L. M., Sullivan, N. L., Cardarelli, L., Septer, A. N., & Gibbs, K. A. (2013). Two independent pathways for self-recognition in *Proteus mirabilis* are linked by type VI-dependent export. *MBio*, *4*(4), e00374-13. https://doi.org/10.1128/mBio.00374-13

Yang, J., Yan, R., Roy, A., Xu, D., Poisson, J., & Zhang, Y. (2015). The I-TASSER Suite: Protein structure and function prediction. *Nature Methods*, *12*(1), 7.

Zepeda-Rivera, M. A., Saak, C. C., & Gibbs, K. A. (2018). A proposed chaperone of the bacterial type VI secretion system functions to constrain a self-identity protein. *J Bacteriol*. https://doi.org/10.1128/JB.00688-17

Zhang, Y. (2008). I-TASSER server for protein 3D structure prediction. *BMC Bioinformatics*, *9*(1), 40.
